# Convergent Evolution of the Antimycobacterial Lasso Peptide Triculamin

**DOI:** 10.1101/2024.12.20.629632

**Authors:** Aske Merrild, Tiziana Svenningsen, Marc G Chevrette, Thomas Tørring

## Abstract

Triculamin is a ribosomally synthesized and post-translationally modified peptide (RiPP) lasso peptide with potent antimycobacterial activity, produced by an unusual, non-canonical biosynthetic gene cluster (BGC). In this study, we elucidate the biosynthetic pathway of triculamin through heterologous expression and show that the biosynthesis proceeds in the presence of a precursor (triA), macrocyclase (triC), and acetyltransferase (triT). Through *in vitro* triT acetylation and bioactivity assays, we show that acetylation functions as a resistance mechanism. Genomic searches of triculamin BGC genes across bacteria show that triculamin is more widely distributed than previously anticipated, as triculamin-like core peptides are found in at least three phyla in contrast to previously described lasso peptides that are typically restricted to one phylum. Triculamin BGCs with both canonical and non-canonical RiPP biosynthetic genes were identified. Two strains containing canonical triculamin-like BGCs were chemically characterized and shown to produce the novel triculamin-like lasso peptides palmamin and gelatinamin, the latter of which appears to have an unprecedented additional ring formation. Detailed phylogenetic investigation of the macrocyclases from triculamin-like BGCs suggests that these molecules are products of convergent evolution. These findings broaden the evolutionary and functional landscape of lasso peptides, revealing their unexpected diversification and cross-phylum distribution.

## Introduction

Ribosomally synthesized and post-translationally modified peptides (RiPPs) are a class of natural products with incredible diversity in structure, biosynthesis, and biological activity.^[1]^ They are typically encoded by precursor peptides located in small biosynthetic gene clusters (BGCs) along with modifying enzymes, transporters, and resistance mechanisms, making them a prime target for genome mining. Several bioinformatic tools, such as antiSMASH^[2]^, RODEO^[3]^, RiPPER^[4]^, RRE-Finder^[5]^, and BAGEL^[6]^, have been developed for this purpose. However, predicting these RiPPs can be challenging due to the small size of the gene clusters and challenges in annotating their precursor peptides. The discovery of new RiPP subclasses in recent years underscores the vast uncharted potential encoded within bacterial genomes.^[7]^

We have previously described the lasso peptide triculamin, an antimycobacterial antibiotic first discovered in 1958, as an unusual member of the lasso peptide family.^[8]^ Not only is it characterized by a large number of cationic amino acids, but it also appears to have a very unusual biosynthesis. The canonical biosynthetic architecture for all other lasso peptides is defined by a precursor peptide (A), a macrocyclase (C) homologous to asparagine synthase, a RiPP recognition element (RRE, B1), and a peptidase (B2). The B1 and B2 are often fused. In addition, the BGC sometimes contains transporters, methyltransferases, and/or other modifying enzymes.^[9–20]^ In contrast, the triculamin BGC, *tri*, appeared to be composed of only the precursor, macrocyclase, and a transporter. More curiously, the triculamin precursor lacked a leader sequence and instead harbored a follower sequence. The macrocyclase was shorter, lacking a domain of unknown function found in other lasso peptide macrocyclases.

Here, we report the heterologous expression of the *tri* BGC in *Streptomyces albus* J1074, elucidate a resistance mechanism, and show that triculamin-like lasso peptides are genetically encoded and produced by several bacterial phyla. Furthermore, we describe the phylogeny of the canonical and non-canonical macrocyclases which reveals their distant evolutionary relationship. A comparison of the characterized and predicted triculamin-like lasso peptides show a high degree of similarity despite their distantly related macrocyclases and BGCs. We therefore hypothesize that the triculamin-like lasso peptides are a rare example of a convergent biosynthesis in natural products, and notably, the first such example for RiPPs, implying that these lasso peptides may play essential functional roles across bacterial phyla.

## Results and Discussion

### Four Genes are Necessary and Sufficient for Triculamin Biosynthesis and Export in Actinobacteria

To investigate the genes required and sufficient for the biosynthesis of triculamin, we amplified the DNA encoding *triACD* from gDNA isolated from *Streptomyces triculaminicus* JCM4242 by PCR and used restriction cloning to insert the putative BGC in the plasmid pL99^[21]^. The plasmid was purified, sequence-verified by whole plasmid sequencing, and transferred into *S. albus* J1074 by *E. coli* ET12567 mediated interspecies conjugation. The extracts of *S. albus* J1074 with the plasmids carrying the BGC and the parental plasmid were compared by LC-MS, MS^2^ and bioactivity testing but showed no indications of triculamin.

Revisiting extracts from *S. triculaminicus*, we discovered several triculamin-like peptides with masses matching an additional Arg (triculamin B) and Arg+Phe (Figure 1a), indicating variable length of the C-terminal tail of the lasso peptide (Table S4). A +42 Da species of all three lasso peptides was also observed, consistent with an acetylation. A similar case was reported by Zong *et. al*., when heterologously expressing the lasso peptide, albusnodin,^[15]^ where the acetylation was performed by the acetyltransferase albT, genomically adjacent to the lasso peptide BGC (*albACB*). They demonstrated that albT was essential for heterologous expression and hypothesized that it likely served as a self-resistance mechanism. Examining the genomic vicinity of the *triACD* BGC, we found a putative acetyltransferase, albeit larger (296 aa vs.178 aa) with low homology to albT (9.4% protein identity/15.4% protein similarity), as well as a gene encoding a transcriptional regulator (H), between triT and *triACD*. We then created the plasmid *pL99-triACDHT* and transferred it into *S. albus* J1074 (Figure 1b). Comparing extracts from *S. albus* J1074 carrying *pL99-triACDHT* or the parental plasmid, *pL99*, we identified triculamin B with a C-terminal Arg and the acetylated triculamin B using LC-MS and MS^2^ (Figure 1c).

**Figure 1.**
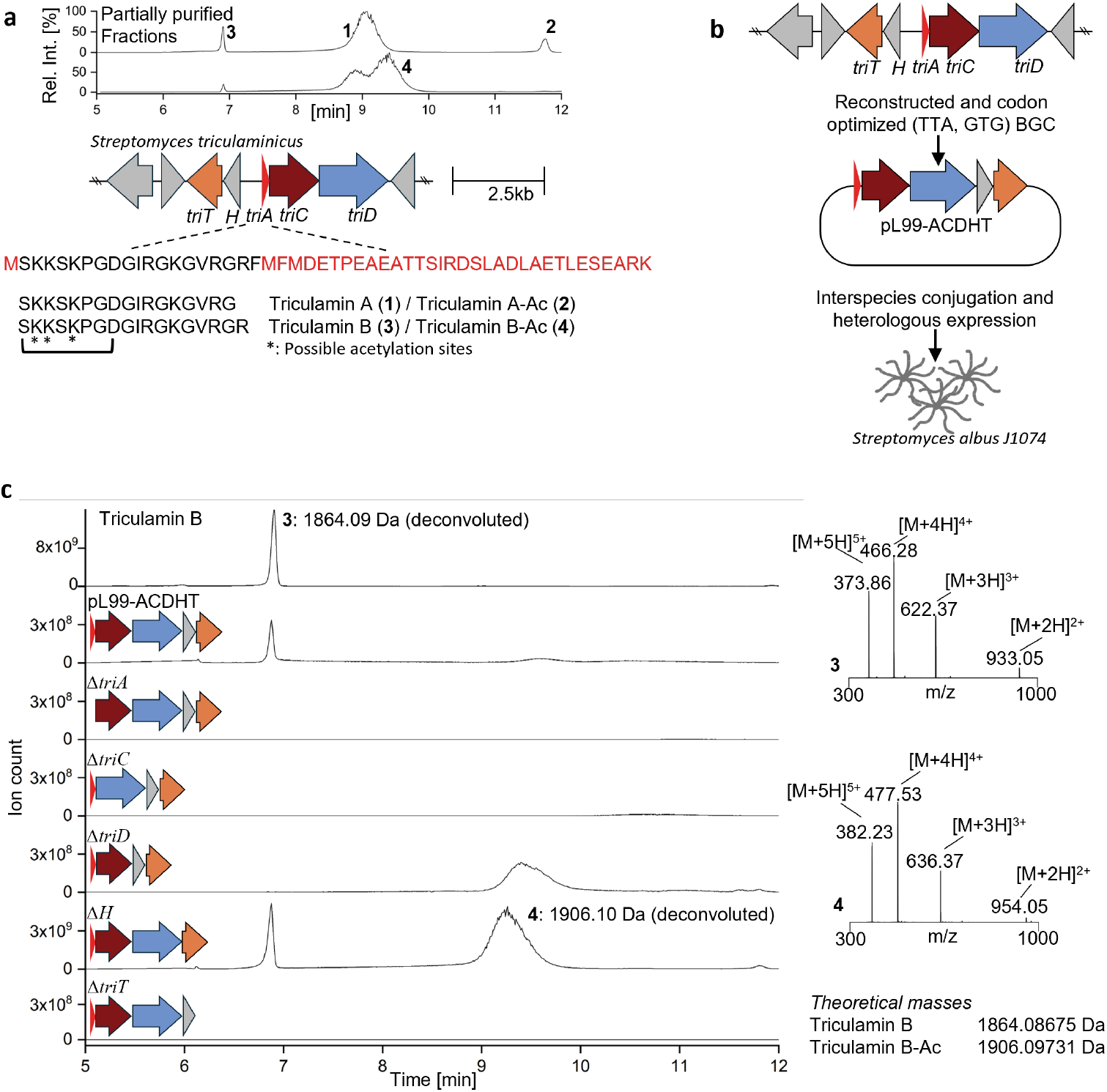
Heterologous production of triculamin in *S. albus*. a) The triculamin BGC and the immediate genetic neighborhood in *S. triculaminicus*. LC-MS traces of extract from *S. triculaminicus* showing peaks corresponding to triculamin A and B as well as their acetylated counterparts. b) The triculamin BGC (including *triT, H, triA, triC*, and *triD*) is cloned into the pL99 plasmid and transferred to *S. albus* J1074 for heterologous production. c) LC-MS deconvoluted traces of individual gene deletions from the pL99 plasmid to investigate the roles of specific genes. MS spectra of triculamin B (3) eluting at 6.9 min and acetylated triculamin B (4) eluting at 9.3 min.

To individually assess the roles of each gene, we created a series of plasmids, each omitting a single gene, transferred them to *S. albus* J1074, and evaluated the extracts using LC-MS and MS^2^. *triA, triC*, and triT were required and sufficient to produce triculamin in *S. albus* J1074 (Figure 1c). The transcriptional regulator H appears to have no role in the heterologous biosynthesis, but could play a regulatory role in the native producer. In the plasmid missing the transporter *triD*, lower amounts are observed, and all observed triculamin is acetylated, suggesting that acetylation occurs while the lasso peptide is trapped in the cytoplasm.

### The N-acetyltransferase triT is a Self-resistance Mechanism

We first hypothesized that acetylation served either as a resistance mechanism or as a post-translational modification occurring prior to macrocyclization, potentially aiding in the prefolding of the precursor into the lasso knot. To validate the role of triT, we synthesized the triT gene codon-optimized for *E. coli*, inserted it in pETM-11 to create a fusion protein with an N-terminal 6xHisTag, and used it to transform *E. coli* BL21 (DE3). The triT-fusion protein was expressed and purified, providing a soluble protein in good yields (∼6 mg/L fermentation broth) (Figure S1). We mixed the triT protein, acetyl-coenzyme A (AcCoA), and triculamin B *in vitro* (Figure 2a) and, following overnight incubation investigated the mixture using both LC-MS, MS^2^ (Figure 2b-c) and bioactivity against *Mycobacterium phlei* and *Mycobacterium smegmatis* (Figure 2d). The results shown in Figure 2, clearly demonstrate that not only is triculamin B fully acetylated (M+42 Da), but it also loses activity against the *Mycobacterium* tested. A closer inspection of the MS^2^ spectra revealed a match consistent with the acetylated lasso peptide observed in the *S. albus* J1074 and *S. triculaminicus* (Figures S9 and S10). Also, it indicates that the acetylation likely happens on one of the three lysine residues (K2, K3, K5) in the lasso macrolactam.

**Figure 2.**
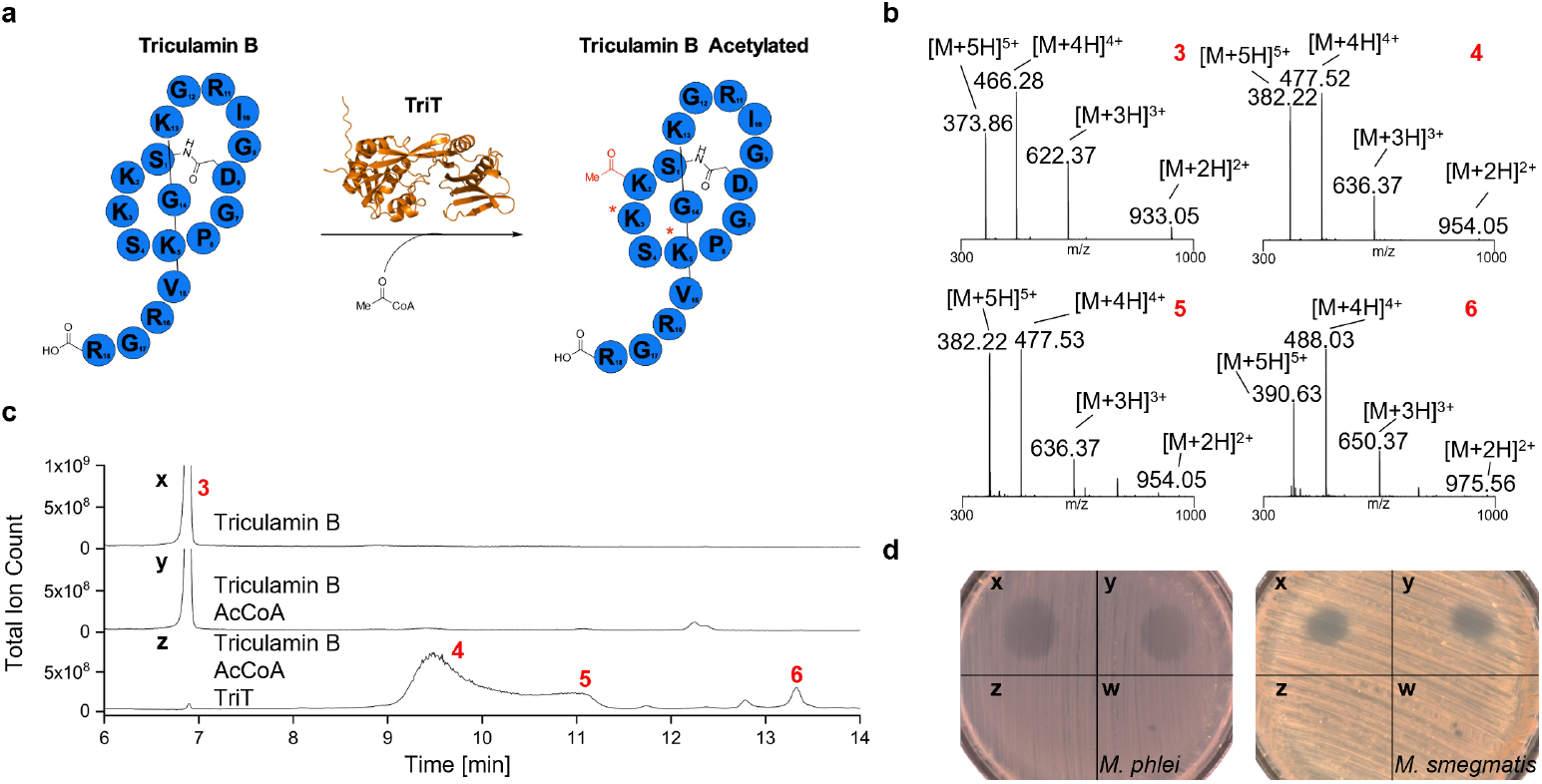
*In vitro* acetylation of triculamin B using purified the acetyltransferase enzyme, triT. a) schematic illustration of the enzymatic reaction. Asterisks 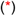 indicate possible acetylation sites. triT is predicted by alpfahold3.^[22]^ b) LC-MS traces of triculamin B (x), triculamin B incubated with AcCoA (y), and triculamin B, AcCoA, and triT (z). c) MS spectra of triculamin B (1) as well as single acetylated (2, 3) and double acetylated (4) species. d) bioactivity from spots of *in vitro* reactions of triculamin B (x), triculamin B incubated with AcCoA (y), triculamin B, AcCoA, and triT (z), and triT and AcCoA(w), tested against *M. phlei* (left) and *M. smegmatis* (right).

### Triculamin-like Peptides are Encoded in both Canonical- and Non-canonical BGCs Across Several Bacterial Phyla

In our efforts to elucidate the essential genes for the heterologous production of triculamin, we noticed a vast number of triculamin-like BGCs found across several bacterial phyla. Some of these clusters were characterized by the unusual follower-containing precursor, the N-truncated macrocyclase, and for most cases, an N-acetyltransferase. Surprisingly, we also identified several canonical lasso peptide BGCs with RREs (B1), peptidases (B2), regular macrocyclase, and precursor peptides with leader sequences. All of these had core sequences nearly identical to triculamin, suggesting that they may follow different biosynthetic strategies yet yield identical lasso peptides. Intriguingly, these appeared across several phyla including Actinomycetota, Bacillota, and Pseudomonadota. To verify the production of triculamin-like peptides in some of the identified bacteria, we selected eight strains for further investigation: *Brevibacillus gelatini, Chitinaciproducens palmae, Paenibacillus tarimensis, Streptomyces alanosinicus, Saccharotrix australiensis, Streptomyces albus* subsp. *chlorinus, Micromonospora inyonensis*, and *Streptomyces kurssanovii*. From both canonical and non-canonical BGCs. These strains were subjected to a One Strain, Many Compounds (OSMAC^[23]^) study (Figures S19 and S20 for results from all but *S. alanosinicus* and *M. inyonensis*) to stimulate the production of the suspected lasso peptide. One strain, *B. gelatini*, successfully produced a triculamin-like peptide under laboratory conditions in addition to the original triculamin strain *S. triculaminicus*.

The major differences in non-canonical and canonical biosynthetic strategies in the identified strains suggest it is unlikely the almost identical predicted lasso peptides of these groups arose from horizontal gene transfer events. This led us to speculate that the potential functional role of triculamin-like lasso peptides could have arisen from convergent evolution. The taxonomic distribution of this lasso peptide family spans three phyla, in stark contrast to most other lasso peptides whose distributions are confined to a single phylum. In the seminal paper describing the development of RODEO,^[3]^ the only cross-phyla example found was propeptin-like peptides that were found in both Actinomycetota and Bacillota. To examine this in more detail, we chose the macrocyclase from experimentally verified producers of triculamin-like peptides; *S. triculaminicus, B. gelatini*, and *C. palmae*, and used these to query against the NCBI non-redundant protein sequences database via protein-BLAST.^[24]^ We manually validated all hits by identifying nearby precursor(s) containing the triculamin-like core peptide (Figure 3, dark purple inner ring - canonical BGCs, light purple inner ring – non-canonical BGC). Macrocyclases of canonical and non-canonical triculamin groups exhibit distinct phylogenetic clustering patterns. These findings underscore the broader evolutionary distribution of triculamin-like peptides and their diversification across bacterial lineages. A full description of how Figure 3 was constructed can be found in the Supporting Information, in addition, a full tree and all Weblogos can be found in Figure S17.

**Figure 3.**
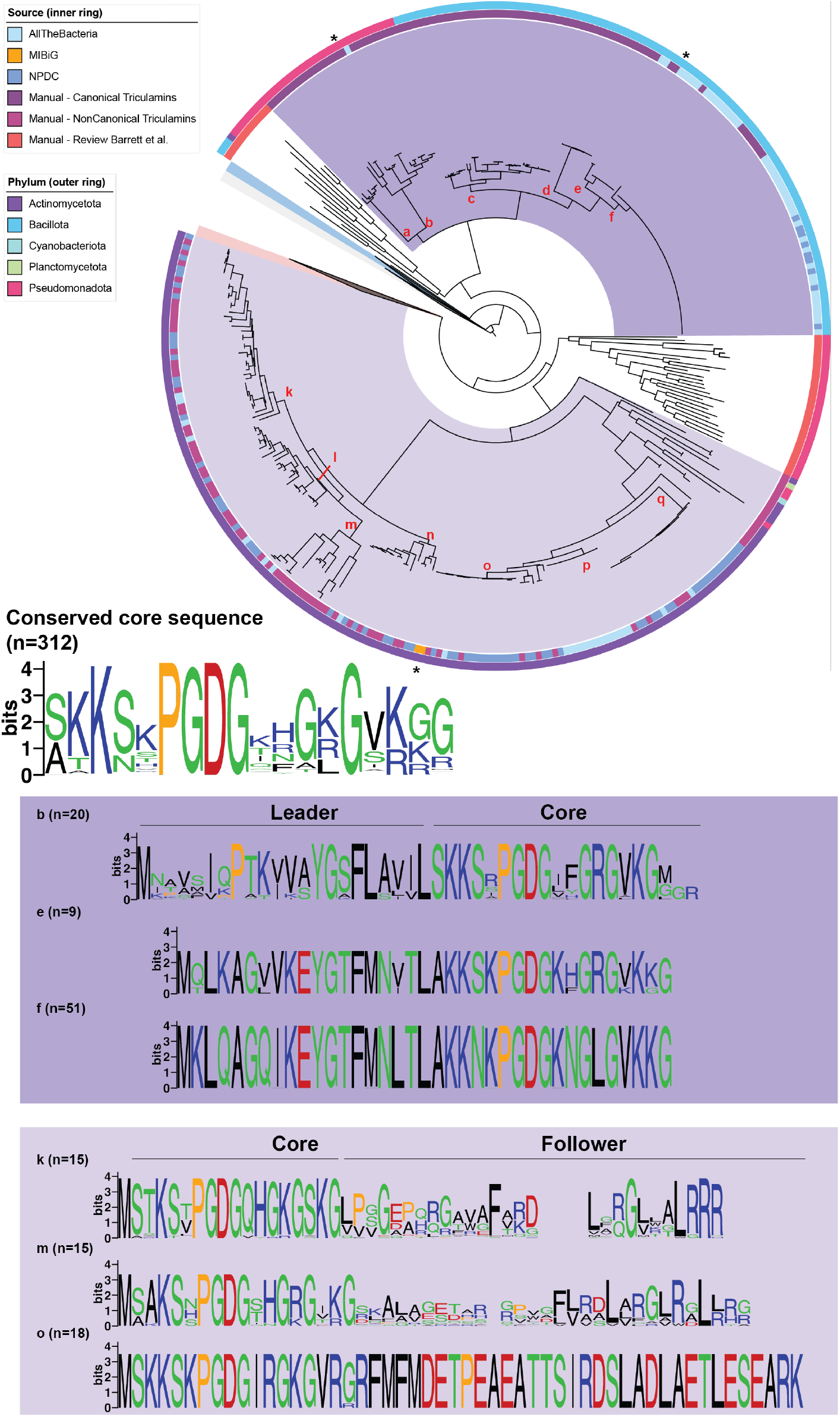
Phylogenetic tree of the macrocyclases. The tree contains three collapsed notes (grey - Asn synthase-like proteins, blue - b-lactam synthases, red – other lasso peptide macrocyclases). Inner background coloring: canonical triculamin-like BGCs = dark purple, non-canonical triculamin-like BGCs = light purple. Weblogo^[44]^ representations of the core sequence for all triculamin-like peptides and several subclades.

The macrocyclase phylogeny reveals that macrocyclases from both the canonical and non-canonical BGCs are found in multiple bacterial phyla. Those from non-canonical BGCs are mainly found in *Actinomycetota (Streptomyces, Micromonospora, Nocardia, Kribella, Lentzea*, etc.), to a lesser extent in *Pseudomonadota* (*Alphaproteobacteria*, genera such as *Ensifer* and *Bradyrhizobium)*, and in a few cases in *Cyanobacteriota* (*Limnospira)* and *Planctomycetota (Rubinisphaera)*. The macrocyclases from canonical BGCs are found in both *Pseudomonadota* and *Bacilliota*. Pseudomonadota is represented by *Betaproteobacteria* within genera such *Trinickia, Paraburkholderia, Burkholderia* and *Gammaproteobacteria* within genera *Xenorhabdus* and *Photorhabdus*. The bacteria *S. triculaminicus, C. palmae* and *B. gelatini* described in detail in this manuscript are marked with an asterisk (*) under clades o, b and e respectively (Figure 3). We also noted that the macrocyclases from other classes of previously characterized lasso peptides, reviewed by Barrett et al.,^[25]^ are distantly related to both macrocyclases from canonical and non-canonical triculamin-like BGCs. A basal Barrett et al. clade (Figure 3, northwest position, orange inner ring) includes macrocyclases such as from the Citrocin^[26]^, Ubonodin^[27]^, and microcin J25^[28]^ BGCs, and the more derived Barrett et al clade (Figure 3, southeastern position, orange inner ring) includes macrocyclases such as from the Capistruin^[29]^, Rubrinodin^[30]^, and Astexin^[31]^ BGC. The light orange collapsed clade is comprised of macrocyclases from other lasso peptides (Supporting Figure S16), including those from the Stlassin^[32]^, Lihuanodin^[10]^, and Deepstreptin^[33]^ BGCs. Interestingly, the southeastern Barrett et al. clade, the collapsed clade containing all other lasso peptides, and the non-canonical triculamin-like clade appear to have been derived from a common ancestor. The more basal canonical triculamin-like clade is sister to the northwestern Barrett et al. clade. The deep branching separating the non-canonical and canonical containing groups suggests independent origins of triculamin-like macrocyclases and convergent or parallel evolution the triculamin-like lasso peptides. The characterized lasso peptides review by Barrett et al. are widely distributed in the extended tree shown in Figure S16, and clearly demonstrates a large diversity in both macrocyclases and core peptides, but surprisingly, we have not been able to identify any non-canonical lasso peptide BGCs encoding precursor other than triculamin-like. We suggest two explanations for this observation: either non-canonical macrocyclases with sufficient diversity to produce distinct lasso peptides have yet to be discovered, or the conserved core sequence facilitates folding, thereby restricting the folding landscape available to non-canonical macrocyclases.^[34,35]^

We compared the precursor peptides of each of the triculamin-like macrocyclase clades (a-f, k-q) and as expected from the diversity in macrocyclases, diversity was observed in the precursors, mostly in the leader and follower sequence regions (Figure 3). The core sequence for all precursors found in both canonical and non-canonical BGCs show an almost invariant PGDG motif centered on the macrolactam-forming aspartate, but also the high number of positively charged amino acids (K, R, H). The leader peptide of canonical clade b stands out as the only one lacking a threonine normally observed in the -2 position. As such, BGC annotation pipelines that require this T (including antiSMASH) fail to identify these precursors. This has also been reported in *Bradymonas sediminis*. ^[36]^ In general, the Y (−17), P (−14), L(−12) and T (−2) motif that is highly prevalent in other leader peptides^[37,38]^ is missing across the canonical clades (a-f). The follower peptides found in the non-canonical BCGs also co-variate with the macrocyclase. The clade o which contains the previously characterized triculamin, shows no diversity in either precursor or macrocyclase. This is in contrast to clades k and m that show conservation in core, but variance in the follower, although they seem to have a central FxRDL motif and a positively charged C-terminal in common. Since the role of the follower peptide is still unknown, we can only hypothesize that this might play a role in binding to the macrocyclase.

Convergent and parallel evolution in natural products and their biosynthesis are striking examples of how bacteria independently evolve similar solutions to ecological challenges. One notable case is the dentigerumycins and gerumycins in *Pseudonocardia* symbionts of fungus growing ants, where structurally similar cyclic depsipeptides are produced despite originating from distinct biosynthetic pathways.^[39]^ This suggests evolutionary pressures in similar niches drive the independent evolution of comparable natural products. ^[40,41]^ Another classic example is found in siderophores, small molecules that chelate iron. While siderophores like enterobactin, pyochelin, and desferrioxamine differ in structure, their shared function highlights convergent evolution shaped by the universal need for iron acquisition in bacteria (ref). Biosynthesis of the clavam ring in β-lactam natural products can be produced by at least four different convergent biosynthetic mechanisms in unrelated bacteria.^[40,42,43]^ These examples underscore how similar environmental pressures may sculpt the evolution of analogous natural products across diverse bacterial lineages.

### Triculamin-like Lasso Peptide from *C. palmae* Heterologously Expressed in *Burkholderia* sp. Ferm-3421

In our attempt to isolate and characterize triculamin-like lasso peptides from canonical lasso BGCs, we screened *Chitinasiproducens palmae*, a member of the Burkholderiaceae family closely related to the genus *Burkholderia*.^[45]^ This particular BGC contains a macrocyclase (palC), an RRE (palB1), a peptidase (palB2), a transporter (palD), and a precursor peptide with a predicted leader sequence in front of the conserved triculamin-like core (Figure 4b). An OSMAC^[23]^ strategy did not produce antimycobacterial compounds, so we opted for a heterologous expression system. An initial attempt to heterologously express the *pal* BGC in *E. coli* BL21(DE3) was unsuccessful (data not shown). Recent advances by Eustáquio and colleagues in heterologous lasso peptide production using the spliceostatin-deficient strain *Burkholderia* sp. FERM-3421 Δfr9, showed great promise as an alternative heterologous host to *E*.*coli*.^[46,47]^ Since *C. palmae* is closely related, we used gDNA to clone the *pal* BGC into expression plasmid pHNF008 and transformed the *Burkholderia* host with pHNF008-palAB1B2CD (Figure 4a). A slightly modified fermentation protocol was designed with a 10-fold reduction in kanamycin concentration and a 10-fold increase in inoculum to mitigate downstream challenges with cation exchange purification caused by high levels of the positively charged aminoglycoside. The chromatogram in Figure 4c clearly shows the production of the triculamin-like lasso peptide, which we term palmamin. The MS and MS^2^ spectra are a clear match to the predicted lasso peptide showing the characteristic fragmentations (b- and y-ions) of the tail.^[48]^ This confirms the successful heterologous expression and demonstrates that distinct biosynthetic machinery found in distantly related bacteria can produce nearly identical lasso peptides.

**Figure 4.**
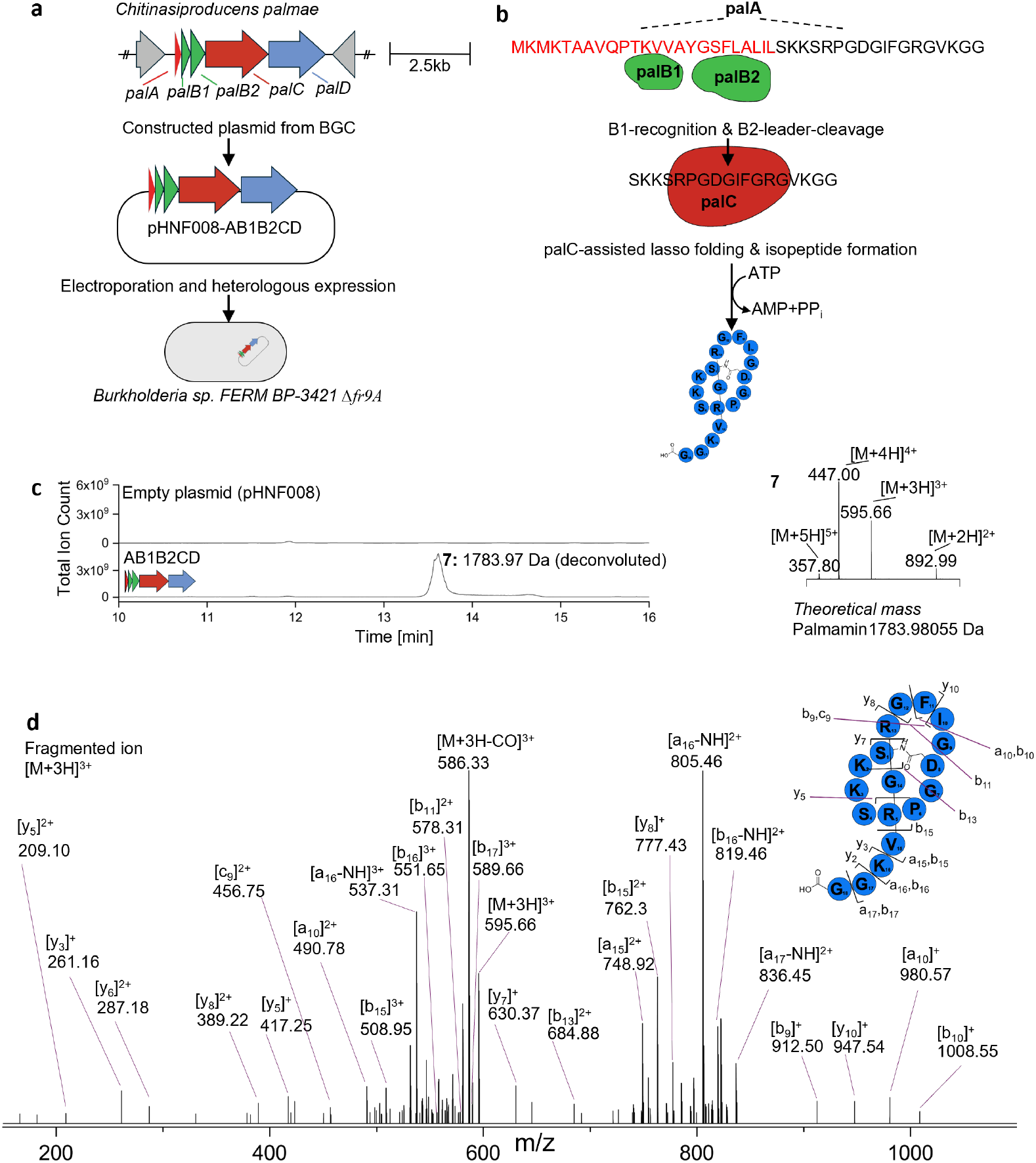
Heterologous expression of the canonical palmamin lasso peptide BGC from *C. palmae*. a) Graphical representation of the *C. palmae* BGC and the workflow for heterologous production in *Burkholderia* sp. FERM BP-3421 Δfr9A. b) Graphical depiction of the hypothesized canonical lasso peptide biosynthesis of palmamin, involving post-translational modification of the precursor peptide by the B1, B2, and C proteins. c) LC-MS traces from the fermentation of *Burkholderia* sp. FERM BP-3421 Δfr9A containing either the empty pHNF0008 plasmid or the plasmid containing the palmamin BGC, showing production of a species with a mass matching the theoretical mass of palmamin. d) MS^2^ fragmentation of the [M+3H]3+ ion of palmamin.

### Isolation of a Proposed Double Cyclized Lasso Peptide Gelatinamin A and its Derivatives from *Brevibacillus gelatini*

As in our approach with *C. palmae* above, we applied an OSMAC strategy to two *Paenibacillaceae* family members, *Brevibacillus gelatini PDF4 (*DSM 100115) and *Paenibacillus tarimensis SA-7-6 (*DSM 19409) (Figures S19). These both encode canonical lasso BGCs with putative triculamin-like lasso peptides (A, B1, B2, D genes). whereas the *B. gelatini* also encodes a clostripain family peptidase (C11-family peptidase) and an N-acetyltransferase (Figure 5a). Unlike with *C. palmae*, here we successfully identified conditions under which *B. gelatini* produced an antimycobacterial compound. This compound exhibited behavior almost identical to triculamin on a cation exchange column, indicating its positively charged nature, and showed a similar elution profile on a C18-polar column (Figure 5b). The masses of the predicted lasso peptide and putative acetylated product could be found in the LC-MS traces and enough for MS^2^ (Figure S11) but were barely observable as minor products. The two major products (**8** and **9** in Figure 5c) had deconvoluted masses of 1741 Da and 1783 Da, respectively. The difference of 42 Da was indicative of an acetylation as seen for triculamin. The discrepancy between the predicted and observed mass is consistent with removal of the C-terminal glycine, followed by an additional dehydration. All expected fragments from the C-terminal tail characteristic of lasso peptides were not observed. The mass difference, lacking tail fragments, and the MS^2^spectra observed in Figure 5d are consistent with macrolactamization between the C-terminus and a lysine residue located in the normal lasso peptide macrolactam. Double fragments indicate it to be on either K2 or K5.^[48,49]^ As in triculamin biosynthesis, the putative gelatinamin BGC contains an N-acetyltransferase, which likely accounts for the observed acetylated products. MS^2^ fragments are consistent with an acetylation of residue K16. Unlike the triculamin BGC, the gelatinamin BGC contains the putative clostripain family protease. Since clostripain is known to catalyze transpeptidase reactions,^[50]^ we hypothesized that the clostripain is responsible for the internal transpeptidase reaction which would yield double the cyclized lasso peptide with glycine removed (see Figure S15).

**Figure 5.**
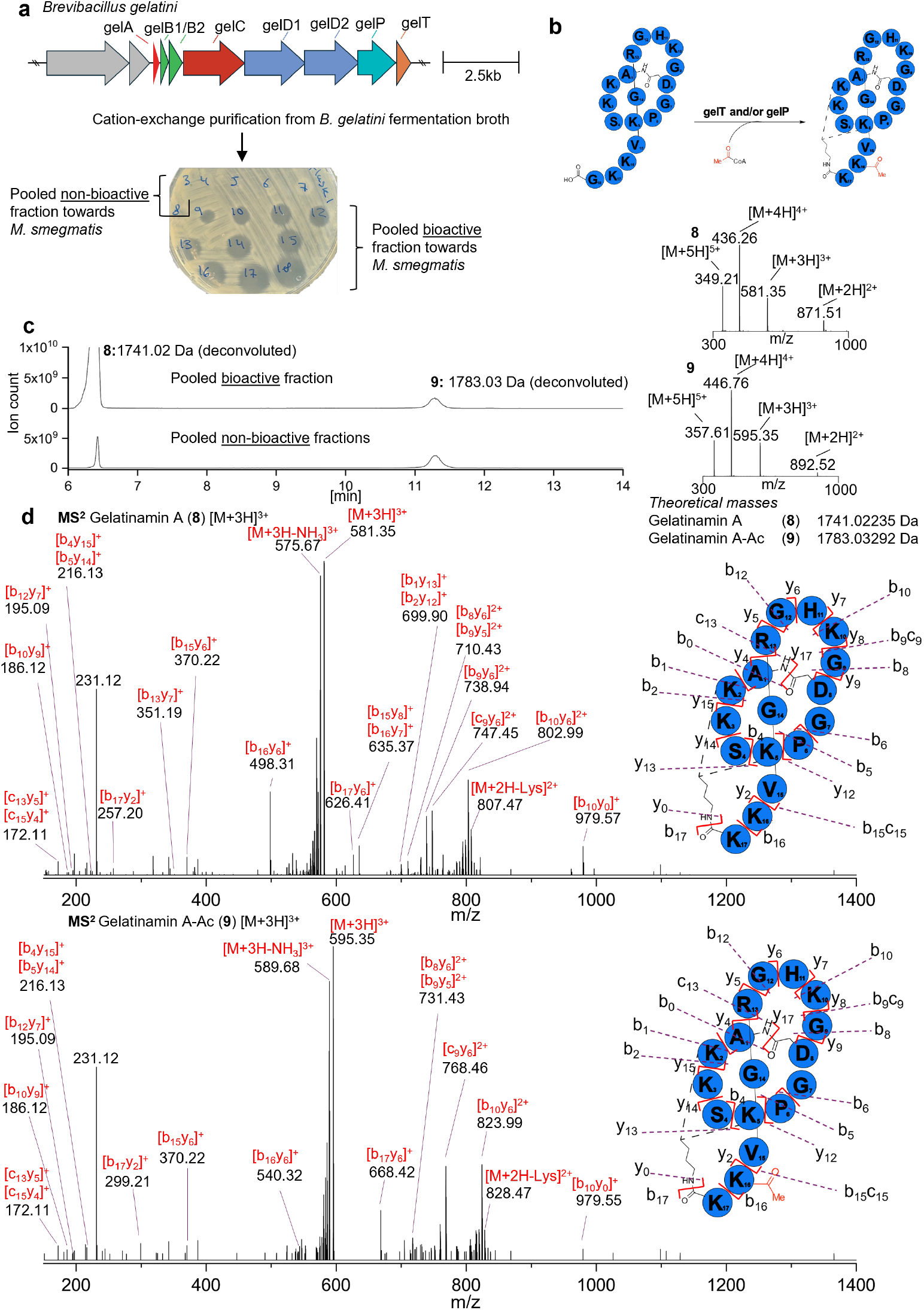
Identification of a lasso peptide containing two macrolactams from *Brevibacillus gelatini*. a) Schematic representation of the gelatinamin BGC from *B. gelatini*. Following fermentation of *B. gelatini*, gelatinamin was isolated using cation exchange chromatography. The elution fractions are shown spotted on a plate with *M. smegmatis*, with fractions 5–8 termed the “non-bioactive” fractions and fractions 9–43 termed the “bioactive fractions” (not all data shown). b) Visualization of the gelatinamin variants identified from *B. gelatini* fermentation, including acetylated and non-acetylated variants, as well as single- and double-dehydrated forms of the lasso peptide. c) LC-MS deconvoluted traces displaying retention times and mass spectra of the pooled non-bioactive and bioactive fractions. The bioactive fractions predominantly consist of non-acetylated gelatinamin, while the non-bioactive fractions contain a mixture of acetylated and non-acetylated variants. Both fractions primarily feature the double-dehydrated form of gelatinamin, with the single-dehydrated form present only in trace amounts. d) MS^2^ spectrum of the gelatinamin A and gelatinamin A-Ac [M+3H]^3+^ peak, chiefly double fragments are observed due to the double cyclization.

A recent preprint by Wright *et al*. describing the lasso peptide lariocidin from *Paenibacillus sp*. M2^[51]^ supports this hypothesis. Coincidently, we have discovered different triculamin-like peptides with canonical BGCs from different members of the *Paenibacillaceae* family, both with the unusual and novel C-terminal to lysine sidechain amine PTM. Through a series of elegant experiments, Wright and colleagues found that lariocidin is an inhibitor of bacterial protein synthesis, and using X-ray crystallography they show the double-cyclized lasso peptide in complex with the small ribosomal subunit in a new site not previously described. Given the overall similarity of the triculamin-like peptide it seems likely that they share the same target and may explain the unusually wide distribution of the BGCs across several bacterial phyla.

### The Acetyltransferase triT Inactivates Gelatinamin via Acetylation

Given the high similarity between triculamin B and gelatinamin A as well as the presence of a predicted acetyltransferase in both BGCs, we hypothesized that acetylation was a general self-resistance mechanism, and that acetylation of gelatinamin A (Figure 6a) also would lead to loss of bioactivity as observed for triculamin A. To test this hypothesis, we evaluated the ability of triT to acetylate gelatinamin A using the same conditions as for triculamin. Our results in Figure 6c demonstrate that triT acetylated gelatinamin A in two positions which completely abolished the bioactivity of gelatinamin A, further supporting its role as a resistance mechanism. Although the sequence identity between *triT* and the native *gelT* is relatively low (44.6%, based on a BLOSUM62 alignment), the close resemblance of their natural substrates likely facilitated the acetylation. However, the results showed that *in vitro* acetylation of gelatinamin A resulted predominantly in a double acetylation of the peptide (Figure 6b). This may be because gelatinamin is naturally acetylated at a different site than triculamin. Analysis of the MS^2^ data suggests that gelatinamin is acetylated on the lysine in the tail (K16), whereas triculamin is most likely acetylated on one of the lysines in the macrolactam (K2,K3, or K5). While the single-acetylated variant is the most abundant form purified from *B. gelatini*, trace amounts of the double-acetylated variant were also detected. Given that the *in vitro* reactions were carried out with a large excess of AcCoA, this surplus likely drove the reaction to saturation, leading to full acetylation at both possible sites. These findings suggest that acetylations could be a widespread strategy for inactivating bioactive lasso peptides, highlighting its potential as a universal resistance mechanism within this RiPP family.

**Figure 6.**
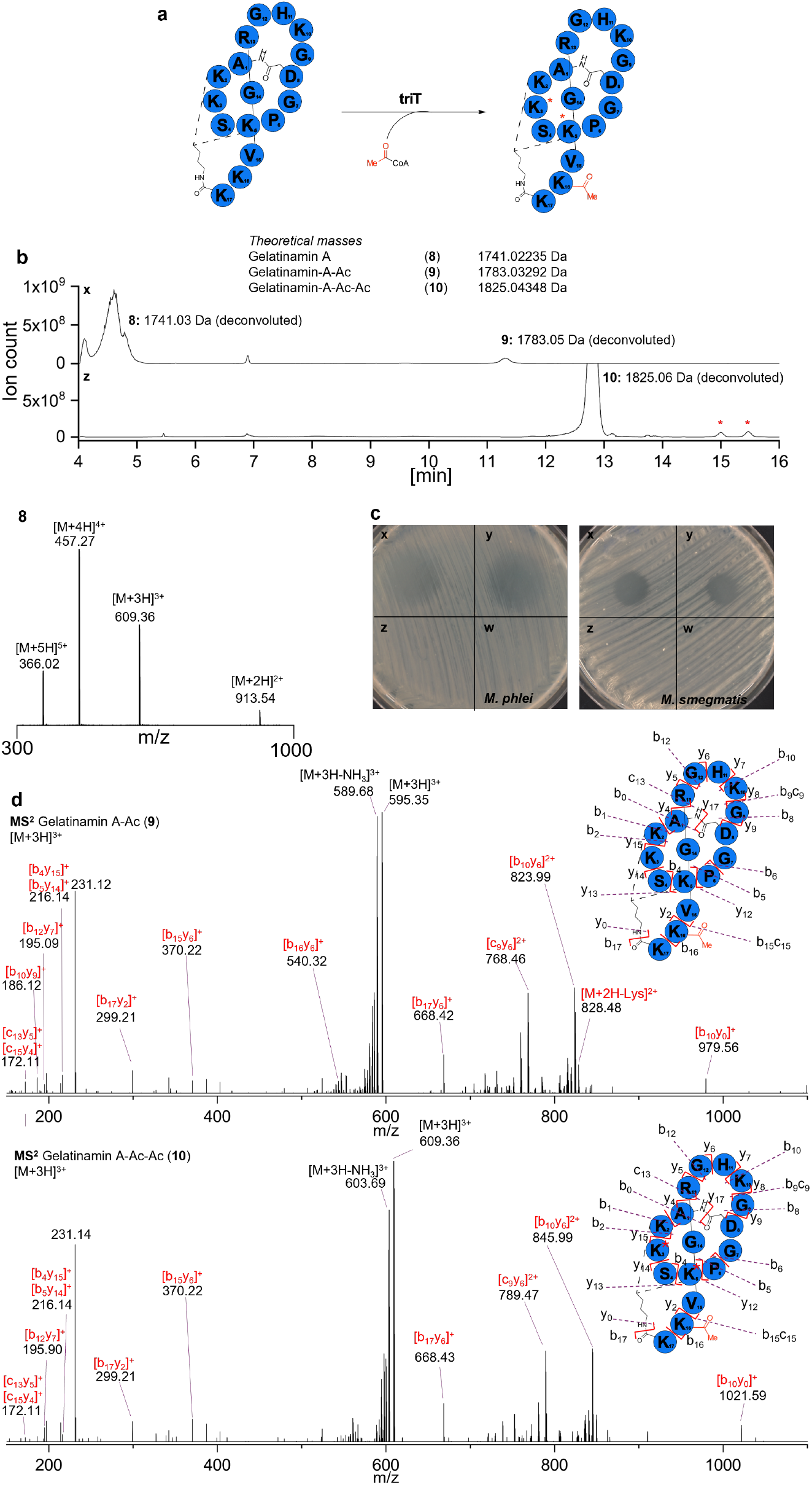
*In vitro* acetylation of gelatinamin using purified triT. a) Schematic of acetylation reaction for single and double acetylation of gelatinamin b) LC-MS traces of gelatinamin (x) and gelatinamin incubated with AcCoA and triT (z), 10 MS spectrum of double acetylated gelatinamin A. * corresponds to peaks with deconvoluted masses fitting triple acetylation of gelatinamin A. c) bioactivity from spots of *in vitro* reactions of gelatinamin (x), gelatinamin incubated with AcCoA (y), gelatinamin, AcCoA, and triT (z), and triT incubated with AcCoA (w). All reactions are tested against *M. smegmatis* and *M. phlei*. d) MS^2^ of gelatinamin A-Ac (9) and double acetylated gelatinamin A-Ac-Ac (10).

## Conclusion

In this manuscript, we demonstrate that the heterologous expression of the triculamin BGC in *S. albus* confirms that four genes—*triA, triC, triD*, and *triT*—are sufficient and necessary for triculamin biosynthesis and export. This represents the first instance of lasso peptide biosynthesis involving a non-canonical BGC, characterized by the absence of an RRE and leader peptidase (B1 and B2), a precursor with a follower sequence, and an N-terminal truncated macrocyclase. The heterologous expression and subsequent purification of triT from *E. coli* enabled *in vitro* acetylation of triculamin. and bioactivity assays, demonstrate acetylation inactivates triculamin, and, together with the necessity of triT in triculamin biosynthesis in *S. albus*, suggests that triT functions as a self-resistance mechanism. Phylogenetic analyses revealed the distribution of both canonical and non-canonical macrocyclases across diverse bacterial phyla harboring triculamin-encoding BGCs. These findings suggest that triculamin represents the first documented example of convergent biosynthesis in a RiPP natural product. Two of the precursors from canonical lasso BGCs were experimentally confirmed to produce triculamin-like lasso peptides (gelatinamin and palmamin), where the gel BGC encodes an additional post-translational modification enzyme, a clostripain-family peptidase. We propose that this enzyme facilitates the internal trans-peptidase reaction required for formation of an additional ring in the lasso peptide gelatinamin A. This hypothesis is supported by MS^2^ spectra and further supported by recent work on lariocidin from the Wright lab.^[51]^ In addition to gelatinamin (A-B), we also confirmed the production of another triculamin-like lasso peptide, palmamin, through heterologous expression in *Burkholderia*. Future research will determine whether this novel post-translational modification and the non-canonical biosynthetic pathway are broadly utilized in lasso peptide biosynthesis or are specific to the triculamin-like peptide family. This work expands the known diversity of lasso peptide biosynthesis, providing new insights into the evolution and functional complexity of non-canonical RiPP pathways. By uncovering both the unique biosynthetic strategies and the potential self-resistance mechanisms, these findings lay the groundwork for exploring triculamin-like peptides as a distinct family with promising biotechnological and therapeutic applications.

## Supporting Information

The authors have cited additional references within the Supporting Information. All raw data has been made available at Open Science Framework (DOI 10.17605/OSF.IO/CZPFE).

## Acknowledgements

We gratefully acknowledge funding from the Carlsberg Foundation (CF22-1239) in the form of a Semper Ardens Accelerate grant [TT] and the Department of Microbiology & Cell Sciences at the University of Florida [MC].

## References

[1] M. Montalbán-López, T. A. Scott, S. Ramesh, I. R. Rahman, A. J. Van Heel, J. H. Viel, V. Bandarian, E. Dittmann, O. Genilloud, Y. Goto, M. J. Grande Burgos, C. Hill, S. Kim, J. Koehnke, J. A. Latham, A. J. Link, B. Martínez, S. K. Nair, Y. Nicolet, S. Rebuffat, H.-G. Sahl, D. Sareen, E. W. Schmidt, L. Schmitt, K. Severinov, R. D. Süssmuth, A. W. Truman, H. Wang, J.-K. Weng, G. P. Van Wezel, Q. Zhang, J. Zhong, J. Piel, D. A. Mitchell, O. P. Kuipers, W. A. Van Der Donk, Nat. Prod. Rep. 2021, 38, 130–239.

[2] K. Blin, S. Shaw, H. E. Augustijn, Z. L. Reitz, F. Biermann, M. Alanjary, A. Fetter, B. R. Terlouw, W. W. Metcalf, E. J. N. Helfrich, G. P. van Wezel, M. H. Medema, T. Weber, Nucleic Acids Research 2023, 51, W46–W50.

[3] J. I. Tietz, C. J. Schwalen, P. S. Patel, T. Maxson, P. M. Blair, H.-C. Tai, U. I. Zakai, D. A. Mitchell, Nat Chem Biol 2017, 13, 470–478.

[4] J. Santos-Aberturas, G. Chandra, L. Frattaruolo, R. Lacret, T. H. Pham, N. M. Vior, T. H. Eyles, A. W. Truman, Nucleic Acids Research 2019, 47, 4624–4637.

[5] A. M. Kloosterman, K. E. Shelton, G. P. Van Wezel, M. H. Medema, D. A. Mitchell, mSystems 2020, 5, e00267–20.

[6] A. J. van Heel, A. de Jong, C. Song, J. H. Viel, J. Kok, O. P. Kuipers, Nucleic Acids Research 2018, 46, W278–W281.

[7] H. Li, W. Ding, Q. Zhang, RSC Chem. Biol. 2024, 5, 90–108.

[8] F. D. Andersen, K. D. Pedersen, D. Wilkens Juhl, T. Mygind, P. Chopin, E. B. Svenningsen, T. B. Poulsen, M. Braad Lund, A. Schramm, C. H. Gotfredsen, T. Tørring, J. Nat. Prod. 2022, 85, 1514–1521.

[9] J. R. Chekan, J. D. Koos, C. Zong, M. O. Maksimov, A. J. Link, S. K. Nair, J. Am. Chem. Soc. 2016, 138, 16452–16458.

[10] L. Cao, M. Beiser, J. D. Koos, M. Orlova, H. E. Elashal, H. V. Schröder, A. J. Link, J. Am. Chem. Soc. 2021, 143, 11690–11702.

[11] C. Zhang, M. R. Seyedsayamdost, ACS Chem. Biol. 2020, 15, 890–894.

[12] S. Zhu, J. D. Hegemann, C. D. Fage, M. Zimmermann, X. Xie, U. Linne, M. A. Marahiel, Journal of Biological Chemistry 2016, 291, 13662– 13678.

[13] S. Zhu, C. D. Fage, J. D. Hegemann, D. Yan, M. A. Marahiel, FEBS Letters 2016, 590, 3323–3334.

[14] L. A. Harris, P. M. B. Saint-Vincent, X. Guo, G. A. Hudson, A. J. DiCaprio, L. Zhu, D. A. Mitchell, ACS Chem. Biol. 2020, 15, 3167–3175.

[15] C. Zong, W. L. Cheung-Lee, H. E. Elashal, M. Raj, A. J. Link, Chem. Commun. 2018, 54, 1339–1342.

[16] E. Gavrish, C. S. Sit, S. Cao, O. Kandror, A. Spoering, A. Peoples, L. Ling, A. Fetterman, D. Hughes, A. Bissell, H. Torrey, T. Akopian, A. Mueller, S. Epstein, A. Goldberg, J. Clardy, K. Lewis, Chemistry & Biology 2014, 21, 509–518.

[17] Z. Feng, Y. Ogasawara, S. Nomura, T. Dairi, ChemBioChem 2018, 19, 2045–2048.

[18] I. Kaweewan, H. Komaki, H. Hemmi, K. Hoshino, T. Hosaka, G. Isokawa, T. Oyoshi, S. Kodani, J Antibiot 2019, 72, 1–7.

[19] T. Zyubko, M. Serebryakova, J. Andreeva, M. Metelev, G. Lippens, S. Dubiley, K. Severinov, Chem. Sci. 2019, 10, 9699–9707.

[20] L. A. Harris, H. Saad, K. E. Shelton, L. Zhu, X. Guo, D. A. Mitchell, Biochemistry 2024, 63, 865–879.

[21] N. Sun, Z.-B. Wang, H.-P. Wu, X.-M. Mao, Y.-Q. Li, Ann Microbiol 2012, 62, 1541–1546.

[22] J. Abramson, J. Adler, J. Dunger, R. Evans, T. Green, A. Pritzel, O. Ronneberger, L. Willmore, A. J. Ballard, J. Bambrick, S. W. Bodenstein, D. A. Evans, C.-C. Hung, M. O’Neill, D. Reiman, K. Tunyasuvunakool, Z. Wu, A. Žemgulytė, E. Arvaniti, C. Beattie, O. Bertolli, A. Bridgland, A. Cherepanov, M. Congreve, A. I. Cowen-Rivers, A. Cowie, M. Figurnov, F. B. Fuchs, H. Gladman, R. Jain, Y. A. Khan, C. M. R. Low, K. Perlin, A. Potapenko, P. Savy, S. Singh, A. Stecula, A. Thillaisundaram, C. Tong, S. Yakneen, E. D. Zhong, M. Zielinski, A. Žídek, V. Bapst, P. Kohli, M. Jaderberg, D. Hassabis, J. M. Jumper, Nature 2024, 630, 493–500.

[23] H. B. Bode, B. Bethe, R. Höfs, A. Zeeck, ChemBioChem 2002, 3, 619.

[24] E. W. Sayers, E. E. Bolton, J. R. Brister, K. Canese, J. Chan, D. C. Comeau, R. Connor, K. Funk, C. Kelly, S. Kim, T. Madej, A. Marchler-Bauer, C. Lanczycki, S. Lathrop, Z. Lu, F. Thibaud-Nissen, T. Murphy, L. Phan, Y. Skripchenko, T. Tse, J. Wang, R. Williams, B. W. Trawick, K. D. Pruitt, S. T. Sherry, Nucleic Acids Research 2022, 50, D20–D26.

[25] S. E. Barrett, D. A. Mitchell, Trends in Genetics 2024, S0168952524001793.

[26] W. L. Cheung-Lee, M. E. Parry, A. Jaramillo Cartagena, S. A. Darst, A. J. Link, Journal of Biological Chemistry 2019, 294, 6822–6830.

[27] W. L. Cheung-Lee, M. E. Parry, C. Zong, A. J. Cartagena, S. A. Darst, N. D. Connell, R. Russo, A. J. Link, ChemBioChem 2020, 21, 1335– 1340.

[28] R. A. Salomón, R. N. Farías, J Bacteriol 1992, 174, 7428–7435.

[29] T. A. Knappe, U. Linne, S. Zirah, S. Rebuffat, X. Xie, M. A. Marahiel, J. Am. Chem. Soc. 2008, 130, 11446–11454.

[30] H. Xiu, M. Wang, C. D. Fage, Y. He, X. Niu, M. Han, F. Li, X. An, H. Fan, L. Song, G. Zheng, S. Zhu, Y. Tong, Biochemistry 2022, 61, 595– 607.

[31] M. Zimmermann, J. D. Hegemann, X. Xie, M. A. Marahiel, Chemistry & Biology 2013, 20, 558–569.

[32] T. Liu, X. Ma, J. Yu, W. Yang, G. Wang, Z. Wang, Y. Ge, J. Song, H. Han, W. Zhang, D. Yang, X. Liu, M. Ma, Chem. Sci. 2021, 12, 12353– 12364.

[33] N. J. Merwin, W. K. Mousa, C. A. Dejong, M. A. Skinnider, M. J. Cannon, H. Li, K. Dial, M. Gunabalasingam, C. Johnston, N. A. Magarvey, Proc. Natl. Acad. Sci. U.S.A. 2020, 117, 371–380.

[34] G. C. A. Da Hora, M. Oh, M. C. Mifflin, L. Digal, A. G. Roberts, J. M. J. Swanson, J. Am. Chem. Soc. 2024, 146, 4444–4454.

[35] S. E. Barrett, S. Yin, P. Jordan, J. K. Brunson, J. Gordon-Nunez, G. Costa Machado Da Cruz, C. Rosario, B. K. Okada, K. Anderson, T. A. Pires, R. Wang, D. Shukla, M. J. Burk, D. A. Mitchell, Nat Chem Biol 2024, DOI 10.1038/s41589-024-01727-w.

[36] Y. Duan, W. Niu, L. Pang, D.-S. Mu, Z.-J. Du, Y. Zhang, X. Bian, G. Zhong, Front. Microbiol. 2023, 14, 1181125.

[37] T. Sumida, S. Dubiley, B. Wilcox, K. Severinov, S. Tagami, ACS Chem. Biol. 2019, 14, 1619–1627.

[38] J. R. Chekan, C. Ongpipattanakul, S. K. Nair, Proc. Natl. Acad. Sci. U.S.A. 2019, 116, 24049–24055.

[39] C. S. Sit, A. C. Ruzzini, E. B. Van Arnam, T. R. Ramadhar, C. R. Currie, J. Clardy, Proc. Natl. Acad. Sci. U.S.A. 2015, 112, 13150–13154.

[40] M. G. Chevrette, C. R. Currie, Journal of Industrial Microbiology and Biotechnology 2019, 46, 257–271.

[41] M. G. Chevrette, K. Gutiérrez-García, N. Selem-Mojica, C. Aguilar-Martínez, A. Yañez-Olvera, H. E. Ramos-Aboites, P. A. Hoskisson, F. Barona-Gómez, Nat. Prod. Rep. 2020, 37, 566–599.

[42] M. A. Fischbach, Current Opinion in Microbiology 2009, 12, 520–527.

[43] F. Barona-Gómez, M. G. Chevrette, P. A. Hoskisson, in Advances in Microbial Physiology, Elsevier, 2023, pp. 309–349.

[44] G. E. Crooks, G. Hon, J.-M. Chandonia, S. E. Brenner, Genome Res. 2004, 14, 1188–1190.

[45] M. Madhaiyan, W.-S. See-Too, R. Ee, V. S. Saravanan, J. S. Wirth, T. H. H. Alex, C. Lin, S.-J. Kim, H.-Y. Weon, S.-W. Kwon, W. B. Whitman, L. Ji, International Journal of Systematic and Evolutionary Microbiology 2020, 70, 2640–2647.

[46] S. Kunakom, A. S. Eustáquio, ACS Synth. Biol. 2020, 9, 241–248.

[47] H. N. Fernandez, A. M. Kretsch, S. Kunakom, A. E. Kadjo, D. A. Mitchell, A. S. Eustáquio, ACS Synth. Biol. 2024, 13, 337–350.

[48] K. Jeanne Dit Fouque, H. Lavanant, S. Zirah, J. D. Hegemann, C. D. Fage, M. A. Marahiel, S. Rebuffat, C. Afonso, Analyst 2018, 143, 1157–1170.

[49] B. Paizs, S. Suhai, Mass Spectrometry Reviews 2005, 24, 508–548.

[50] S. Yagisawa, S. Watanabe, T. Takaoka, H. Azuma, Biochemical Journal 1990, 266, 771–775.

[51] G. Wright, M. Jangra, D. Travin, E. Aleksandrova, M. Kaur, L. Darwish, K. Koteva, D. Klepacki, W. Wang, M. Tiffany, A. Sokaribo, B. Coombes, N. Vázquez-Laslop, Y. Polikanov, A. Mankin. preprint 2024, DOI:10.21203/rs.3.rs-5058118/v1.

