## Supporting Information for "Convergent Evolution of the Antimycobacterial Lasso Peptide Triculamin"

[b] Assist. Prof. M. G. Chevrette

Department of Microbiology & Cell sciences

University of Florida, Gainesville, Florida, USA

[c] University of Florida Genetics Institute

University of Florida, Gainesville, Florida, USA

Supporting information for this article is given via a link at the end of the document.

Figure S17 ... 23
3.7 Observed Variants of Lasso Peptides ... 23
Figure S18 ... 23
Table S4 ... 23
3.8 OSMAC Strategy of Canonical BGC Containing Strains ... 24
Figure S19 ... 24
Figure S20 ... 24
References ... 25

### 1. Data Accessibility

We made files that may be of interest available and can be found online at open science framework (OSF) (DOI 10.17605/OSF.IO/CZPFE). This includes cloning files (constructed plasmids, sequencing results, primers). HPLC-MS Raw data (Orbitrap Exploris™ 120 Mass Spectrometer). Manually annotated and predicted lasso peptide BGCs triculamin-analogs (139 from canonical lasso-BGC structure, 174 from non-canonical lasso-BGC structure) and the BGCs of 80 experimentally characterized lasso peptides.[1]

### 2. Methods

#### 2.1 General

Synthetic DNA: Synthetic DNA primers were obtained from Integrated DNA Technologies (IDT) (found at OSF). The t*riT* gene was codon-optimized for *E. coli* K12 expression and synthesized by IDT. The gene was designed with 20-nucleotide overhangs at the N- and C-termini to facilitate cloning into the pETM11 vector (see under Heterologous expression of TriT in the Open Science Framework).

Reagents: Phusion Plus polymerase and dNTPs were supplied by Thermo Scientific. DreamTaq and DreamTaq Green polymerase master mix were also obtained from Thermo Scientific.

Reagents: DNase I from bovine pancreas was supplied by Sigma-Aldrich. Acetyl-CoA trilithium salt (Cat. No. 10101893001) was also obtained from Sigma-Aldrich. A stock of AcCoA was prepared by dissolving the powder in dissolved in phosphate buffer (pH 7.4) at a concentration of 3 mg/mL. Aliquots were prepared and stored at -70°C.

DNA Assembly: NEBuilder® HiFi DNA Assembly Master Mix was obtained from New England Biolabs, and 5X In-Fusion Snap Assembly Master Mix was supplied by TaKaRa Bio.

Enzymes: All FastDigest enzymes used for molecular cloning were supplied by Thermo Scientific. Golden Gate Enzymes used were BsaI-HF®v2 and T4 DNA Ligase provided by New England Biolabs.

Plasmid and DNA Purification: The GeneJET Plasmid Miniprep Kit was used for plasmid purification and obtained from Thermo Fisher. GeneJET Genomic DNA Purification Kit was used for genomic DNA isolation according to manufacturer's protocol obtained from Thermo Fisher. The Monarch PCR & DNA Cleanup Kit and the Monarch® DNA Gel Extraction Kit (Cat. No. T1120) were supplied by New England Biolabs.

Gel Electrophoresis: UltraPure™ Agarose (Invitrogen) was used for gel analysis. SYBR Safe DNA Gel Stain was employed alongside the 1 kb GeneRuler (Cat. No. SM1333). For samples not pre-stained in the mastermix, 6X DNA Loading Dye (Cat. No. R0611) was used to stain the samples.

SDS-PAGE: SDS-PAGE of proteins was conducted using 4–12% Criterion™ XT Bis-Tris Protein Gels with 1XMOPS buffer. Samples were prepared using XT Sample Buffer (Cat. No. 1610791) and analyzed with the PageRuler™ Prestained Protein Ladder (Cat. No. 26616) as a marker.

Protein concentrations: Vivaspin 6 centrifugal concentrator (10 kDa MWCO, 5 mL) was supplied by Sigma-Aldrich and was used for *TriT*.

Dialysis: Spectra/Por® 3 dialysis membrane tubing was utilized for imidazole removal following His-Tag purification. For *TriT*, a MWCO of 12-14 kDa was employed and supplied by Spectrum Labs (spectrumlabs.com).

Competent Cells: Chemically competent *E. coli* DH5α cells were used for plasmid maintenance, while BL21(DE3) cells were employed for protein production and purification. Both competent cell types were prepared in-house and used for transformation following established protocols.[2]

Sanger Sequencing: All Sanger sequencing, including whole plasmid and primer-based supreme seq sequencing, was conducted by Eurofins Genomics. The results of all sequencing can be found on the OSF. LC-$MS^2$ Analysis: LC-MS, $MS^2$ analyses were performed using a Thermo Scientific Orbitrap Exploris™ 120 Mass Spectrometer, coupled with a Thermo Scientific SII LC system. The MS ionization source used electron spray ionization. Data from mass spectrometry were processed using Freestyle (Thermo Fisher) software.

Recipes of all media used in the study can be found in Table S1, and all buffers used can be found in Table S2.

**Table S1.** List of media recipes.

| Name | Composition |
|---|---|
| Mueller Hinton broth (MHB) | Dextrose 1.0 g/L, tryptone 5.0 g/L, yeast extract 2.5 g/L |
| Mueller Hinton Agar (MHA) | Dextrose 1.0 g/L, tryptone 5.0 g/L, yeast extract 2.5 g/L, agar 9.0 g/L |
| ISP2 Broth | Yeast extract 4.0 g/L, malt extract 10.0 g/L, glucose 4.0 g/L, pH 7.4 |
| ISP2 Agar | Yeast extract 4.0 g/L, malt extract 10.0 g/L, glucose 4.0 g/L, agar 15 g/L, pH 7.4 |
| LB Broth | Tryptone 10.0 g/L, yeast extract 5.0 g/L, NaCl 10.0 g/L |
| LB Agar | Tryptone 10.0 g/L, yeast extract 5.0 g/L, NaCl 10.0 g/L, agar 15.0 g/L |
| Triculamin production media (TPM) | Glucose 22.0 g/L, yeast extract 2.5 g/L, NH4NO3 4.0 g/L, CaCO3 2.0 g/L, NaCl 2.0 g/L |
| Tryptic soy broth (TSB) | Peptone from casein 17.0 g/L, peptone from soymeal 3.0 g/L, glucose monohydrate 2.5 g/L, NaCl 5.0 g/L, di-Potassium hydrogen phosphate 2.5 g/L |
| Tryptic soy agar (TSA) | Peptone from casein 17.0 g/L, peptone from soymeal 3.0 g/L, glucose monohydrate 2.5 g/L, NaCl 5.0 g/L, di-Potassium hydrogen phosphate 2.5 g/L, agar 15 g/L |
| MS agar | D-mannitol 20.0 g/L 20.0 g/L, fat-reduced soy flour 20.0 g/L, agar 20.0 g/L, pH 8 After autoclavation add pre-autoclaved $MgCl_2$ to a final concentration of 10 mM |
| YEME medium without sucrose (YEME-WF) | 3.0 g/L Yeast extract, 3.0 g/L malt extract, 5.0 g/L peptone, 10.0 g/L glucose after autoclavation add pre-autoclaved $MgCl_2$ to final concentration 5 mM |
| ISP4 media | 10.0g/L soluble starch, 1.0 g/L K2HPO4, 1.0 g/L MgSO4·7H2O, 1.0 g/L NaCl, 2.0 g/L (NH4)2SO4, 2.0 g/L CaCO3, 1 mg/L FeSO4, 1 mg/L MnCl2, 1 mg/L ZnSO4 |
| ATCC media | 10.0 g/L glucose, 20.0 g/L soluble starch, 5.0 g/L yeast extract, 5.0 g/L N-Z-Amine type A, 1.0 g/L CaCO3, pH 7.3 |
| altMS media | 10.0 g/L mannitol, 10.0 g/L soy flour, 10 g/L malt extract |

**Table S2.** List of buffer concentrations.

| Name | Composition |
|---|---|
| Lysis Buffer | 20 mM Phosphate Buffer (pH 7.4), 200 mM NaCl, 0.2 mg/ml lysozyme, 20 µg/ml DNAse, 1 mM $MgCl_2$ |
| Dialysis buffer | 20 mM Phosphate Buffer (pH 7.4), 200 mM NaCl |
| Cation exchange buffer A | 50 mM HEPES (pH 8) |
| Cation exchange buffer B | 50 mM HEPES (pH 8), 1M NaCl |
| LB Broth | Tryptone 10.0 g/L, yeast extract 5.0 g/L, NaCl 10.0 g/L |
| LB Agar | Tryptone 10.0 g/L, yeast extract 5.0 g/L, NaCl 10.0 g/L, agar 15.0 g/L |
| His-tag purification buffer A | 20 mM Phosphate (pH 7.4), 500 mM NaCl, 10 mM Imidazole |
| His-tag purification buffer B | 20 mM Phosphate (pH 7.4), 500 mM NaCl, 500 mM Imidazole |

#### 2.2 Microbial Strains and Material

The native producer of triculamin, *Streptomyces triculaminicus* (JCM 4242) was acquired from the Japan Collection of Microorganisms, *Brevibacillus Gelatini PDF4* (DSMZ100115), *Paenibacillus tarimensis SA-7-6* (DSM19409), *Chitinasiproducens palmae JS23* (DSMZ 27307) were acquired from the German Collection of Microorganisms and Cell Cultures GmbH (DSMZ). *Streptomyces kurssanovii* 10294 (NRRL B-3366), *Saccharothrix australiensis* LL-BM 782Ce82 (NRRL 11239), *Streptomyces albus subsp. chlorinus* (NRRL B-24108), *Micromonospora inyonensis* INA (NRRL 3292), and *Streptomyces alanosinicus* (B-3627) were acquired from the ARS Culture Collection (NRRL). *Streptomyces albus J1074* used for heterologous production, and *Escherichia coli ET12567* used for interspecies conjugation were kindly provided by Tilmann Weber. *Burkholderia sp.* FERM BP-3421 ΔFr9 was kindly provided by Alessandra S Eustáquio.[3]

#### 2.3 Production and Purification of Triculamin Variants from *Streptomyces triculaminicus* JCM4242

*S. triculaminicus* was cultured on ISP2 agar plates and incubated (28°C, 7 days) to promote sporulation. After incubation, sterile saline water (0.9% NaCl) was added to the plate, and a sterile cotton swab was used to gently dislodge the spores. The spore suspension was used to inoculate six baffled Erlenmeyer flasks (500 mL) containing ISP2 (100 mL). These starter cultures were incubated at (28°C, 180 rpm, 5 days), before being used as inoculum in six baffled Erlenmeyer flasks (5000 mL) containing triculamin production medium (TPM, 2 L). The production cultures were incubated (28°C, 220 rpm, 14 days). After fermentation, cells were harvested by centrifugation (8000 rpm, 20 min, 4°C), and the supernatant was filtered through a coffee filter. The entire 12-liter volume of filtered supernatant was incubated overnight with 1 kg of HP20 resin (28°C, 40 rpm). The following morning, the resin was collected by filtration and washed with water. Triculamin was eluted from the resins using 100% methanol. The aqueous methanol was reduced by rotary evaporation and followed by freeze-drying. While this purification strategy was employed based on prior success[4], we now recognize that the solid-phase extraction step could have been bypassed. Instead, the supernatant could have been used directly for cation exchange chromatography, as described below. This adjustment would likely have increased the overall yield.

#### 2.4 Construction of Phylogenetic Tree of Macrocyclases from Triculamin-like BGC

The macrocyclases from the experimentally verified producers; *S. triculaminicus*, *B. geletini*, and *C. palmae* were used to query against the NCBI non-redundant protein sequences database via BLAST.[5] We manually validated all hits by identifying nearby precursor(s) containing the triculamin-like core peptide (Figure 3 marked "manual", either dark purple inner ring - canonical BGCs, light purple inner ring – non-canonical BGC). Additionally, we initiated an iterative search of MIBiG v4.0rc1[6] with diamond v2.0.15.153[7]. Briefly, an initial query containing i) macrocyclases of all the acquired strains (*B. gelatini , M. inyonensis, S. australiensis, C. palmae, S. alanosinicus, S. albus subsp chlorinus, S. triculaminicus, P. tarimensis* and *S. kurssanovii* ), ii) the albusnodin/ikarugamycin family macrocyclase (accession WP_040248810.1), and iii) the ancestral asparagine synthetase B (accession AAA23498.1) searched MIBiG (--ultra-sensitive -k 0 -b 20 -c 1 -id 25). The hits were then used as queries in a subsequent MIBiG search (--ultra-sensitive -k 0 -b 20 -c 1 -id 35). This was repeated for 8 iterations, where new hits were no longer found. All macrocyclases from the lassopeptide-containing clade were used as diamond queries (--ultra-sensitive -k 0 -b 20 -c 1 --id 60 --query-cover 60) of the AllTheBacteria collection[8] (v0.1; light blue inner ring) and the Natural Product Discovery Center (NPDC) collection[9] (accessed 2024-09-10; dark blue inner ring). We also included a large subset of previously characterized macrocyclases reviewed by Barrett et al.[1] (orange inner ring). Identified macrocyclases were aligned mafft v7.508[10] and the resulting multiple sequence alignment was trimmed using trimAl v1.4.rev15[11] with the -automated1 option, which applies an automatic algorithm to optimize alignment quality by removing poorly aligned regions. Phylogenetic trees were inferred using FastTree v2.1.11[12] with the following parameters: the LG model of amino acid substitution (-lg), accommodating rate heterogeneity across sites using a discrete gamma model (-gamma) with 60 rate categories (-cat 60). Support values were estimated from 1,000 bootstrap replicates (-boot 1000). The macrocyclases in canonical and non-canonical triculamin group were all manually evaluated for the presence of a precursor sequence.

#### 2.5 Alignment of lasso peptides from canonical and non-canonical triculamin BGCs

The genomic vicinity of the canonical and non-canonical macrocyclases (found in 2.4) was manually evaluated for the presence of a precursor peptide. All except a few (three macrocyclases) contained a core sequence like triculamin. The three examples missing a precursor was likely due to small contig size.

The precursors were grouped according to their macrocyclase clades (see Figure S16) and alignments were performed with the clustalX multiple protein alignment.[13] Alignment files were evaluated for wrongful annotation of precursor start codons (actinomycetes commonly use GTG which wrongfully is annotated Val instead of Met). An alignment was made with all the precursors, and the alignment was trimmed only to include the core sequences (leader sequence, follower sequence and core reduced as few positions in the loop and tail region in few cases allow insertions/deletions).

Alignments were also made for each of the clades represented in Figure 3. The WebLogos were prepared as described by Crooks GE et al.[14]

##### 2.6 Heterologous Expression of triT in *E. coli* BL21(DE3)

For expression of triT, the pETM-11 vector was used. The whole plasmid sequencing of the empty plasmid can be found at OSF. The pETM-11 vector was linearized by PCR and fused with the triT insert using In-Fusion® cloning with a 1:3 molar ratio (vector:insert). Chemically competent *E. coli* DH5α cells were transformed with the In-Fusion® reaction mixture following standard protocol and plated on selective media LB agar containing kanamycin (50 µg/mL). Following overnight incubation single colonies were cultured in LB media with kanamycin (50 µg/mL) (37 °C, 140 rpm, overnight). Plasmids were purified (Miniprep Kit K0503) and submitted to whole-plasmid sequencing. Chemically competent *E. coli* BL21(DE3) was transformed with sequence-verified plasmids for protein expression. Single colonies from the transformation plates were picked and used to inoculate baffled Erlenmeyer flask containing LB broth (100 mL) of LB media with kanamycin (50 µg/mL) and incubated (37°C, 140 rpm, overnight). The overnight cultures were used to inoculate large-scale production in LB media (3 x 2L with baffled Erlenmeyer flask) containing kanamycin (50 µg/mL) at a 1:100 dilution ratio. The flasks were incubated (37 °C, 140 rpm) and the optical density (OD600) was monitored until it reached 0.4–0.8. Once the desired OD600 was reached, the cultures were cooled (ice bath, 15 min), production induced with IPTG (final conc. 0.5 mM), and incubated (18°C, 140 rpm, overnight). Following incubation, the cells were harvested by centrifugation (8000 rpm, 20 min, 4°C) and the pellet stored (-20°C) until further use.

##### 2.7 Purification of triT

Frozen cell pellets from the expression of triT were thawed, weighed, and resuspended in lysis buffer (Table S2) at a ratio of 4 mL buffer per gram of cell pellet. The resuspended cells were stirred using a magnetic stirrer at room temperature for 30 minutes. Following enzymatic lysis, the lysates were subjected to sonication using a MS73 probe (60% amplitude, 5 min, 1 sec on, 1 sec off). This sonication program was repeated once. Following sonication, the lysates were centrifuged (12000 rpm, 20 min, 4 °C). The supernatants were then filtered (0.4 µm) before application on the ÄKTA™ start system.

Purification was performed on an ÄKTA™ start chromatography system using a 5 mL HisTrap FF Crude column (11000458). For purification a flow rate of 5 mL/min was used. The complete chromatography program, including wash and elution gradients, is detailed in Table S3. For triT purification, UV absorbance was monitored, and fractions with significant UV signals were collected for SDS-PAGE analysis to confirm the presence of the protein (Figure S1).

Fractions containing the target protein were pooled and dialyzed against dialysis buffer (Table S2) to remove the imidazole. The dialyzed proteins were concentrated to a final concentration of 1 mg/mL using centrifugal filter devices Vivaspin 6 centrifugal filters (10 kDa MWCO) from Sigma-Aldrich. The concentrated protein was stored at -70 °C.

**Table S3.** AKTA program for his-tag purification of triT

| **Column** | **5 mL HisTrap FF crude** | |
|---|---|---|
| **Flowrate** | **5 mL/min** | |
| **Steps** | **Action** | **Duration (CV)** |
| **1** | Equilibrating column in water | 2 |
| **2** | Equilibration of column in buffer A | 5 |
| **3** | Sample application | - |
| **4** | Column wash – wash unbound sample (buffer A) | 10 |
| **5** | Elution, gradient (0% to 25% buffer B) | 10 |
| **6** | Elution, isocratic (100% buffer B) | 10 |
| **7** | Equilibration in buffer A | 5 |

##### 2.8 SDS-PAGE of triT

SDS-PAGE was performed to analyze the elution fractions obtained from his-tag purification and to identify fractions containing the target protein. Based on the UV absorbance profile recorded by the ÄKTA system (Supporting Figure S1), selected fractions were tested. To prepare samples, 5 µL of 4XT sample buffer (#1610791) was mixed with 15 µL of the elution fraction. The mixtures were boiled (95 °C, 15 min). Subsequently, 15 µL of each prepared sample was loaded onto a 4–12% Criterion™ XT Bis-Tris Protein Gel. A molecular weight marker (#26616) was also loaded for reference. The gel was run for 60 minutes at 140 V using MOPS running buffer. After electrophoresis, the gel was stained with READYBLUE® protein gel stain (Sigma-Aldrich) to visualize the proteins.

##### 2.9 Enzymatic Acetylation Reaction of Triculamin and Gelatinamin with triT

The purified lasso peptides, and the acetyltransferase (triT) were thawed on ice and mixed at a 3:1 molar ratio along with 32.25 mM AcCoA (final concentration). The reaction mixture was incubated (18 h, 28 °C, 180 rpm). The reactions were terminated by freezing the vials (-20 °C) and storing them (-20 °C). The following control reactions were also prepared: (1) gelatinamin/triculamin in water (2) gelatinamin/triculamin with AcCoA (3) triT with AcCoA, and finally (4) gelatinamin/triculamin with triT and AcCoA. These controls were treated under identical conditions to the main reaction.

Part of the reaction was subject to HPLC-MS analysis: The debris was removed by centrifugation (14000 rpm, 5 min), and the supernatant was analyzed by Orbitrap MS and $MS^2$ with 0.1 % formic acid (FA) in water as running solvent and 0.1 % FA in MeCN as elution solvent.

The rest of the mixture was used for bioactivity screening as described below.

##### 2.10 Plate Based Bioactivity Screening of *in vitro* Reactions

Bioactivity tests were conducted against *Mycobacteriuam smegmatis* and *Mycobacteriuam phlei* on MHA plates. Frozen aliquots stored at -70°C were thawed and plated on fresh MHA plates, then incubated (37°C, overnight). For all tests, a sterile cotton swab was dipped into a culture of either *M. smegmatis* or *M. phlei*, and a fine, even smear of culture was spread across a fresh MHA plate. Drops (20 µL) of the test substances were then spotted onto the smeared plates. Plates were incubated (37°C, overnight) and inspected the following morning. To ensure a clear inhibition zone boundary, the plates were left at room temperature for a few additional days before final evaluation and imaging. Plates were imaged using ProtoCOL 3HD (Synbiosis, Cambridge, UK) with an exposure time of 440 ms. All results from bioactivity screening of *in vitro* reactions can be found in Figures S2, S3, S4 and S5.

##### 2.11 Construction of Plasmids for Heterologous Expression in *Streptomyces albus* J1074

All necessary information for cloning, plasmids and sequencing results can be found on the OSF database.

Scheme S1a illustrates the overall workflow of pL99 derived plasmid preparation[15] For all PCR-amplifications the Phusion Plus polymerase was used according to the provided protocol (with GC-enhancer used). Restriction enzymes were FastDigest and used according to manufacturer protocol. Initially, *triACD* was PCR-amplified (primers triA_F and triD_R) from genomic *S. triculaminicus* introducing NdeI and HindIII sites followed by gel electrophoresis and gel extraction. pL99 and the *triACD* fragments were digested with NdeI and HindIII, followed by gel electrophoresis and gel extraction. Purified fragments were ligated with T4 DNA ligase (Thermo Fisher) according to the provided protocol. 100 µL In-house-made chemically competent *E. coli* DH5α were transformed with 10 µL ligation mixture. Following overnight growth on LB agar plates supplemented with apramycin (50 µg/mL), colonies were picked for growth in liquid LB-apramycin (50 µg/mL) and grown over night. The cultures were used for purification of plasmid pL99-triACD and sent to whole plasmid sequencing. From genomic DNA *triHT* was PCR amplified (primers triT_F and triH_R) introducing a BglII and HindIII site, followed by gel electrophoresis and gel extraction. Next, *triHT* fragment and pL99-triACD were digested using BglII and HindIII, followed by gel electrophoresis and gel extraction. Purified fragments were ligated with T4 DNA ligase (Thermo Fisher) according to the provided protocol. 100 µL In-house-made chemically competent *E. coli* DH5α were transformed with 10 µL ligation mixture. Following overnight growth on LB-apramycin (50 µg/mL) agar plates, colonies were picked for growth in liquid LB-apramycin (50 µg/mL) and grown over night. The cultures were used for purification of plasmid pL99-"*triACDHT*" and sent to whole plasmid sequencing. Next, undesired codons GTG and TTA were mutated into ATG and CTG. First, 5 separate PCR reactions (primers 4.1-4.5), using pL99-"*triACDHT*" as a template, were performed. Fragments were evaluated by gel electrophoresis and purified by gel extraction. The purified fragments were assembled using NEBuilder using provided protocol. 100 µL In-house-made chemically competent *E. coli* DH5α were transformed with 10 µL NEBuilder mixture. Following overnight growth on LB-apramycin (50 µg/mL) agar plates, colonies were picked for growth in liquid LB-apramycin (50 µg/mL) and grown over night. The cultures were used for purification of plasmid pL99-triACDHT and sent to whole plasmid sequencing. To create the knock-out variants of the triBGC the pL99-triACDHT was used as a template. As the large plasmid (12.2 Kb) and high GC-content (66%) gave rise to troublesome PCRs, we found that knocking out a gene by a single PCR was inefficient. Thus, each knock-out was achieved by assembling two PCR fragments with NEBuilder. The construction of each fragment follows the outlined method in Scheme S1a.

##### 2.12 Construction of Plasmid for Heterologous Expression in *Burkholderia sp. FERM BP-3421 ∆fr9*

All necessary information for cloning, plasmids and sequencing results can be found on database OSF database.

In contrast to the pL99 plasmid, pHNF008 plasmid is readily available for Golden Gate cloning (except for one present BsaI site). Thus, primers to both integrate the palAB1B2CD and remove the undesired BsaI site was designed using the online tool NEBridge Golden Gate (New Englands Biolabs). The plasmid fragments were PCR amplified (primers 6.1F/6.1R and 6.2F/6.2R) using pHNF008 as template. The palAB1B2CD was PCR amplified (6.3F/6.3R) from *C. palmae* genomic DNA. Gel electrophoresis was performed for all three reactions and fragments were gel-extracted. Golden Gate cloning was performed according to manufacturer manual from New England Biolabs and in-house-made chemically competent *E. coli* DH5α was transformed and selected on LB-kanamycin (50 µg/mL) agar. Following overnight growth on LB-kanamycin (50 µg/mL) agar plates, colonies were picked for growth in liquid LB-kanamycin (50 µg/mL) and grown over night. The cultures were used for purification of plasmid pHNF008-palAB1B2CD and sent to whole plasmid sequencing

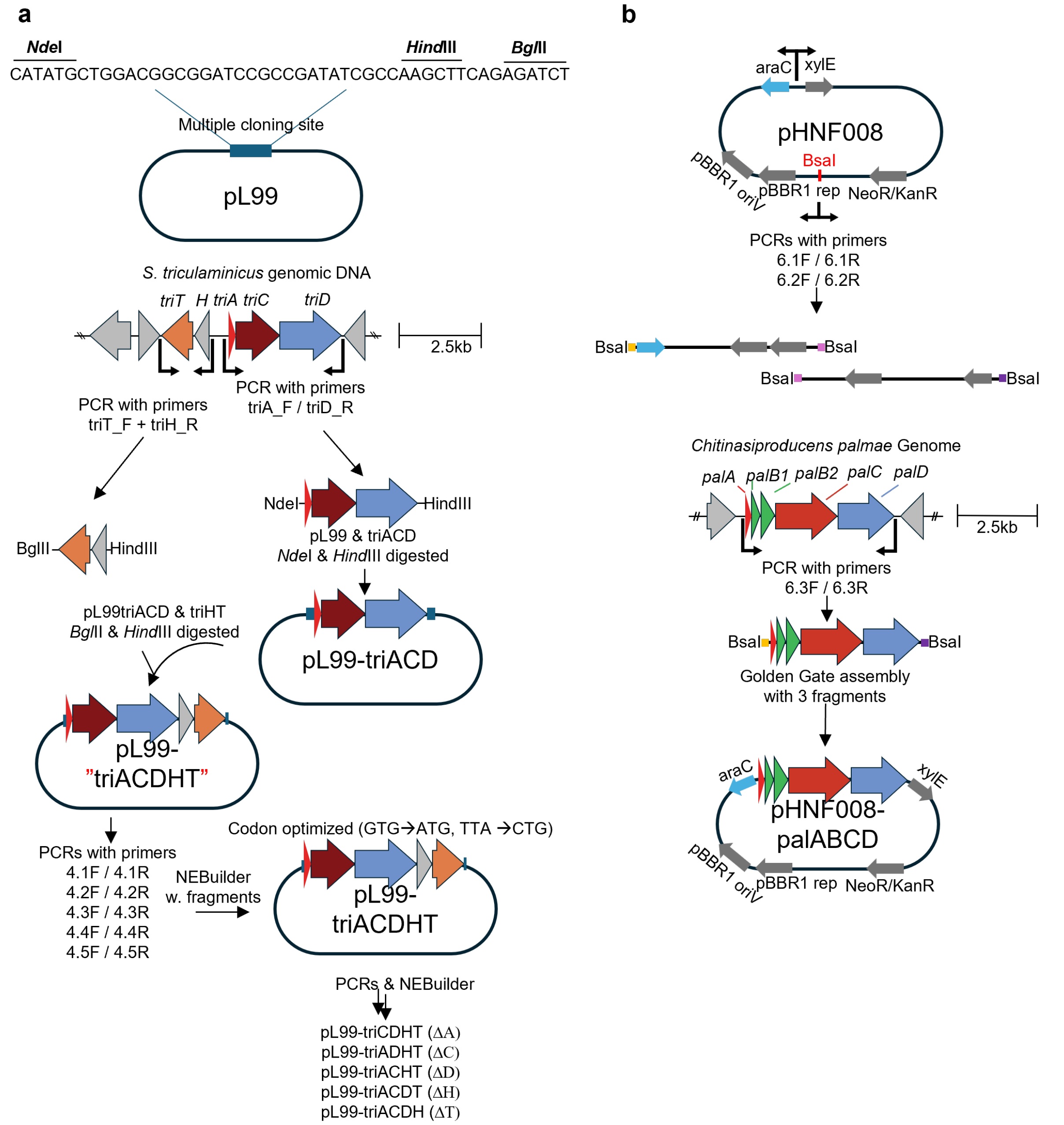

**Scheme S1**. The overall workflow of cloning procedures for construction of heterologous expression plasmids. a) construction of plasmids for heterologous expression in *S. albus* J1074. The procedure uses both restriction-cloning and NEBuilder for assembly. b) construction of the plasmid for heterologous expression in *B. sp FERM-3421 ΔFr9*. The plasmid was assembled with a Golden Gate cloning strategy, while removing one undesired BsaI site from the pHNF008 plasmid.

##### 2.13 Heterologous Production of Triculamin in *Streptomyces albus J1074*

Expression plasmids were transformed according to a previously described protocol.[16] In short, DNA-methylation deficient *E. coli* ET12567 was transformed with the plasmid of interest and harvested. Next, harvested spores of *S. albus* J1074 were cultivated together with transformed *E. coli* to achieve interspecies conjugation. The mixture was negatively selected towards apramycin and nalidixic acid and positive colonies were plated on ISP2 supplemented with apramycin (50 µg/mL) agar plates. Single colonies were carefully spread on MS agar supplemented with apramycin (50 µg/mL) and grown for 3-5 days until fully sporulated. Each spore plate was harvested by 2x10 mL YEME medium without sucrose (YEME-WS) and 10 mL spore suspension was used to inoculate 40 mL YEME-WS apramycin (final concentration 50 µg/mL). The culture was grown (28°C, 180 rpm, baffled Erlenmeyer flask) for one day, followed by induction with ε-caprolactam (final concentration 0.5 % w/v). After 5 days the fermentation broth was collected. We have previously shown that triculamin is highly heat-stable, thus lysis of cells was achieved by 30 min at 95°C followed by centrifugation (12000 g, 45 min). The supernatant was collected. When the transporter (triD) is present triculamin accumulates in the supernatant and thus lysis of the cells is unnecessary.

##### 2.14 Heterologous Production of Palmamin in *Burkholderia sp. FERM BP-3421 ∆fr9*

Heterologous expression from *Burkholderia sp. FERM BP-3421 ∆fr9* followed the method described by Fernandez et al.[3] In short, the *Burkholderia* strain was electroporated with the desired plasmid and selected towards kanamycin. A single colony from LB agar supplemented with kanamycin (0.5 mg/mL) was used to inoculate 10 mL LB kanamycin (0.5 mg/mL) and grown 1-2 days in seed medium (28°C, 180 rpm, baffled Erlenmeyer flask, until OD600 of 5-8). As the high concentration of kanamycin interfered with our downstream purification steps, we reduced the concentration in the production medium to 50 µg/mL. To compensate for the reduced antibiotic concentration, we increased the inoculum (final OD600 of 0.1 instead of 0.01). Therefore, lasso peptide production was achieved by inoculation to 2S4G medium containing kanamycin (50 µg/mL) and L-arabinose (100 mM) and cultivated for 5 days. Cells and supernatants were harvested by centrifugation (12000g, 45min). Palmamin was found in the supernatant.

##### 2.15 Production and Purification of Lasso Peptide from *Brevibacillus gelatini* PDF4

A single colony of *B. gelatini* was picked from an agar plate of TSA agar and used to inoculate 200 mL TSB. The starter culture was incubated (37 °C, 180 rpm, 48 h). The starter culture was used to inoculate 6x2L of TSB media in baffled Erlenmeyer flasks in a 1:100 ratio and incubated (37 °C, 140 rpm, 5 days). Following fermentation, cells were harvested (4 °C, 12000 rpm, 20 min). The supernatant was buffered with HEPES (50 mM, pH 8) and filtered through a coffee filter at least 3 times before storing (4 °C).

Purification was performed by cation exchange on an ÄKTA™ start chromatography system using a 5 mL HiTrap SP XL column (17-5160-01). For the purification, a flow rate of 5 mL/min was used. The complete chromatography program, including wash and elution gradients, is detailed below. The purification was performed over 6 runs, with 2L buffers supernatant for each run. For each run all fractions eluted were tested for bioactivity against *M. smegmatis* (data not shown). We hypothesized that the acetylated fractions would elute before the non-acetylated ones, and to confirm this we tested some fractions on MALDI-TOF MS (data not shown). As anticipated, acetylated started to elute around fraction 5, continuing till fraction 9-11, with a small shift between the 6 purification rounds.

All fractions with bioactivity were pooled as well as the none-bioactive and dialyzed against dd$H_2O$ to remove all salts using 500 MWCO dialysis bags.

##### 2.16 Preparation and Cation Exchange Purification of Lasso Peptides

In our previous exploration of triculamin, a comprehensive purification strategy was applied due to limited knowledge of the physicochemical properties of triculamin. Thus, with the knowledge of triculamin, we designed a cation exchange purification for the highly positively charged lasso peptides. Extracts containing lasso peptides from either native bacterial producers or heterologous expression were buffered (HEPES 50 mM, pH 8) to reduce unspecific unbinding on column and filtered through (0.45 µm). Different methods were designed, depending on the purpose of cation exchange. Method 1 was designed to concentrate and partially purify the heterologously expressed lasso peptides prior to HPLC-MS analysis, whereas method 2 was used for the purification of lasso peptides from native producers. Both methods used mobile phase A (50 mM HEPES, pH 8), mobile phase B (50 mM HEPES, 1M NaCl, pH 8) and flow of 5 mL/min. Prior to sample application, the cation exchange column (HiTrap SP XL, 5mL, Cytiva) was equilibrated with 5 CV A, followed by application of sample and wash of unbound sample with 2-5 CV A. Method 1 eluted lasso peptides by 100% B for 5 CVs and fractions were collected in 2 mL aliquots. Method 2 utilized a two-step gradient. First, gradient was 0-10% B over 10 CV. Secondly, an elution gradient of 10-100% B over 20 CV collecting fraction of 2 mL. UV280 was measured but gave no signal at the time of lasso peptide elution. The methods designed were run on an Äkta GO system. Sample volumes of 50-200mL were used with method 1 and sample volumes from 100 mL to 2L were loaded with method 2. No bioactivity against *M. smegmatis* was detected in the wash fraction indicating no loss of lasso peptide. Lasso peptide containing fractions were determined by bioactivity against *M. Smegmatis* as described in previous section or by MALDI-TOF MS as described below.

##### 2.17 HPLC-$MS^2$ Method

The HPLC- $MS^2$ was performed using an Orbitrap Exploris 120 Mass spectrometer (Thermo Fisher) coupled with a Thermo Scientific SII LC system. A Luna Omega Polar C18 column (5µm, 100Å, 250x4.6mm) was used. Mobile phase A (100% $H_2O$, 0.1% FA) and mobile phase B (100% Acetonitrile, 0.1% FA) were used. The temperature of the column was kept at 30°C, flow was kept at 0.5 mL/min, all samples were filtered (0.22 µm) and the injection volume was 10 µL. The following method was used: 0-2 min 5% B, from 2-13 min a linear gradient from 5% to 25% B, from 13-18 min a linear gradient from 25% to 95% B. At last, the column was washed and re-equilibrated. The Orbitrap Exploris 120 MS received eluate from 4-5 min to 18 min. The ESI-MS conditions were as follows; Spray voltage static at 3500 V, gas mode set at static with sheath gas at 40 AU, auxiliary gas 8 AU and sweep gas at 1 AU. The ion transfer tube temperature was set at 300°C and vaporizer temperature at 320°C. The polarity was positive, colision induced dissociation (CID) was used for fragmentation, Orbitrap resolution set to 120000, Scan range 200-1000 m/z and refraction lens set to 70%. Data dependent $MS^2$ was acquired. Data analysis was performed with FreeStyle software (chromatograms, MS, deconvolution, $MS^2$).

##### 2.18 MALDI-TOF MS

MALDI-TOF MS was used to evaluate the fractions from the cation exchange purification. Fractions were pooled depending on MALDI-TOF spectrum and bioactivity against *M. Smegmatis*. 20 µL of each fraction was acidified (5µL of 10% trifluoracetic acid) and desalted by in-house made disposable Zip-tip columns. The material used for Zip-tip columns were C18AR spec-disc (Agilent). A small piece of C18 material was cut using a blunt syringe-needle and inserted into a 20 µL pipette tip. A 1mL pipette-tip fit into the 20µL tip, which allowed the use of a 1mL pipette for eluting the column. First, the Zip-tip was flushed with 20 µL acetonitrile, next it was flushed with 20µL dd$H_2O$ 0.1% TFA followed by application and elution of the acidified sample. The column was washed with 20µL dd$H_2O$ 0.1% TFA and eluted using 70% acetonitrile 0.1% TFA. The eluate was dried at 50°C and resuspended in 2-4 µL 70% acetonitrile 0.1% TFA saturated in matrix (α-Cyano-4-hydroxycinnamic acid). 1µL sample was spotted onto MTP 384 polished steel plate (Bruker). The instrument used was Bruker an Autoflex MALDI-TOF MS from Bruker, the method used was the default reverse positive method with settings scan range 700-3500 m/z.

##### 2.19 OSMAC of Ordered Strains with Triculamin-like BGCs

In addition, we obtained three strains containing canonical triculamin-like BGCs: *Paenibacillus tarimensis* SA-7-6, *Brevibacillus gelatini* PDF4, and *Chitinasiproducens palmae* strain JS23. These strains were assessed for bioactivity against *M. smegmatis* using patch/patch assays on TSA agar plates.

The three strains were also cultivated in liquid TSA medium and incubated (37°C, 140 rpm, 4 days). After incubation, the fermentation broth was separated by centrifugation (12,000 rpm, 20 min), and both the supernatant and pellet were heat-treated (98°C, 15 min) prior to use in drop assays against *M. smegmatis*.

To further enhance and stimulate production, an OSMAC (One Strain Many Compounds) approach was employed, varying production media (M9 and TSA), fermentation time (3 to 7 days), and oxygen availability (90 to 180 rpm). Fermentations were assessed for bioactivity following the method described in “Plate based bioactivity screening of *in vitro* reactions”. Results can be found in Supporting Figures S19 and S20.

#### 3. Supporting Results and Discussion

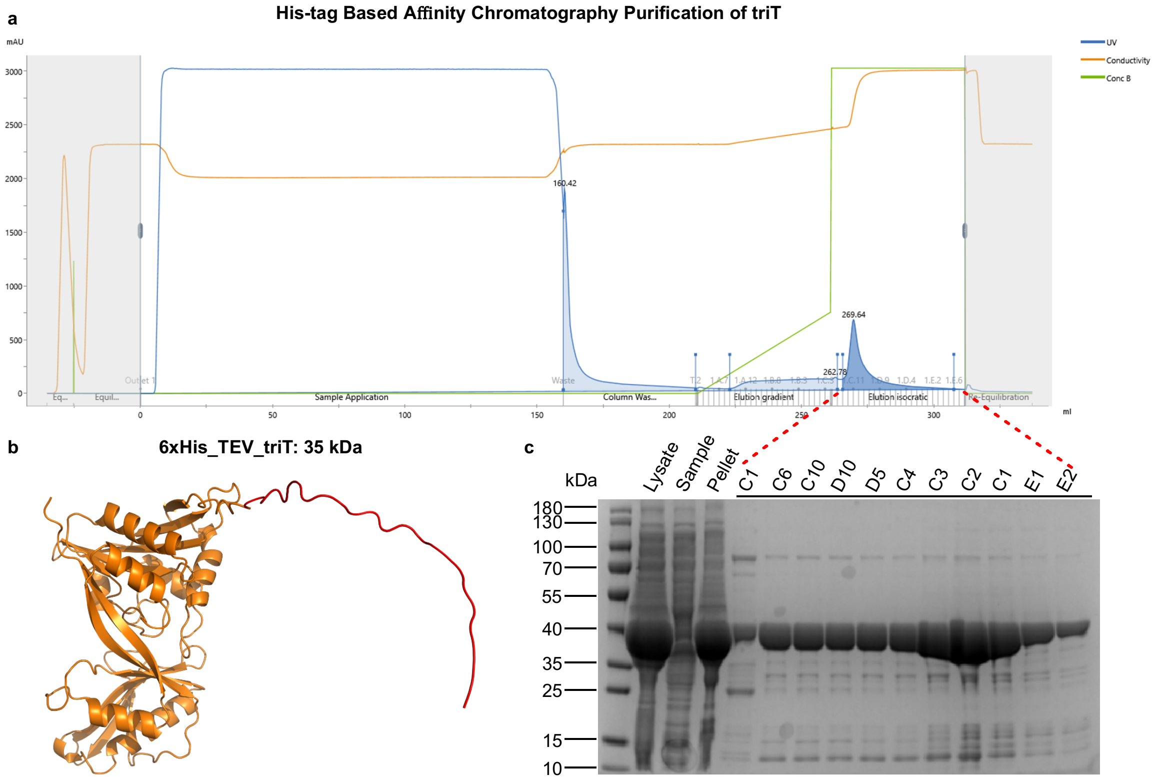

**Figure S1**. Purification of triT. a) Chromatogram from his-tag purification performed on the ÄKTA go system, illustrating all phases, including equilibration, sample application, elution (gradient and isocratic), and finally re-equilibration. A distinct UV-peak is observed shortly after the switch to isocratic elution. b) AlphaFold 3 prediction of the His-tag and triT fusion protein, with a theoretical mass of ~35 kDa. The red region of the protein represents the His-tag and TEV site, and the orange region triT. c) SDS-PAGE analysis of control samples and elution fractions. A prominent band at ~35 kDa is visible in all elution fractions, indicating successful expression and purification of triT.

##### 3.1 Bioactivity Results

Purified triT was utilized *in vitro* to acetylate triculamin B/gelatinamin B by combining triculamin B/gelatinamin B, triT, and AcCoA. The reaction mixtures, along with control samples, were incubated and subsequently tested for bioactivity against *M. smegmatis* and *M. phlei*.

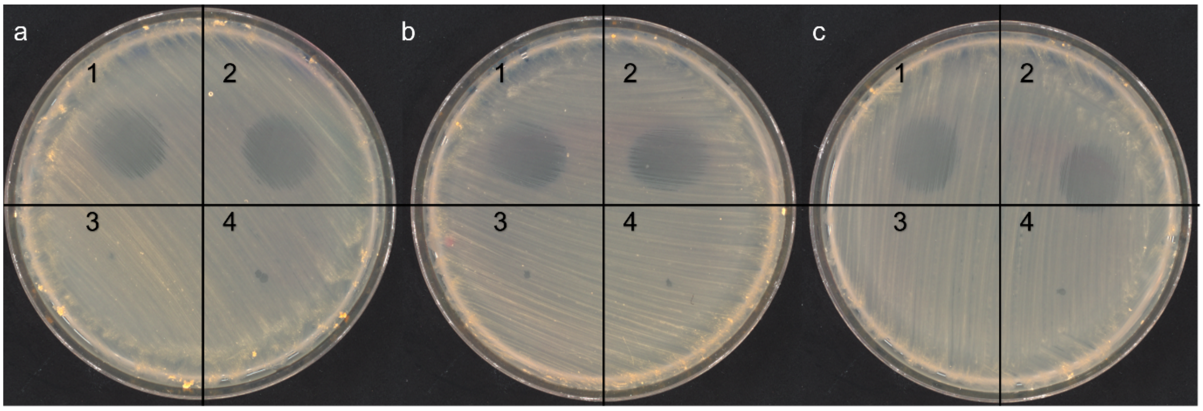

**Figure S2.** Bioactivity of testing of *in vitro* reactions; 1) Triculamin B. 2) Triculamin B incubated with AcCoA 3) Triculamin B incubated with AcCoA and triT 4) triT incubated with AcCoA. All samples are tested in triplicate (a, b and c) against *M. phlei*.

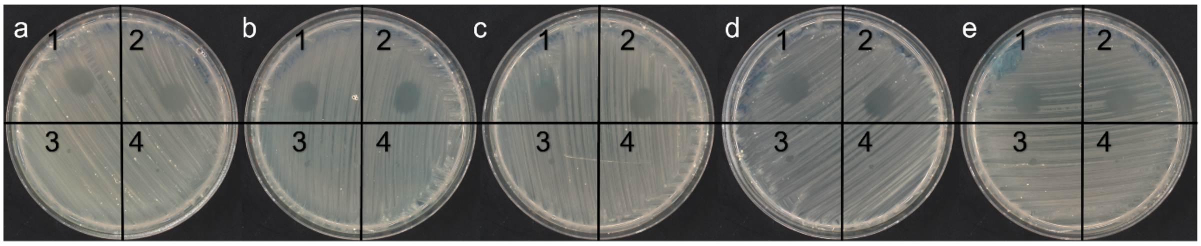

**Figure S3.** Bioactivity of testing of *in vitro* reactions; 1) Triculamin B 2) Triculamin B incubated with AcCoA 3) Triculamin B incubated with AcCoA and triT 4) triT incubated with AcCoA. All samples are tested in quintuplicates (a, b, c, d and e) against *M. smegmatis*.

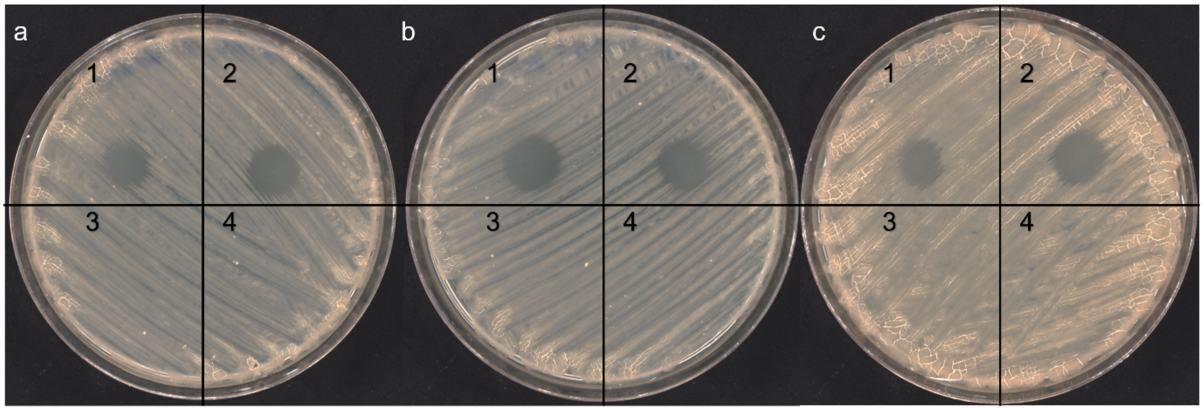

**Figure S4.** Bioactivity of testing of *in vitro* reactions; 1) Gelatinamin 2) Gelatinamin incubated with AcCoA 3) Gelatinamin incubated with AcCoA and triT 4) triT incubated with AcCoA. All samples are tested in triplicate (a, b and c) against *M. smegmatis*.

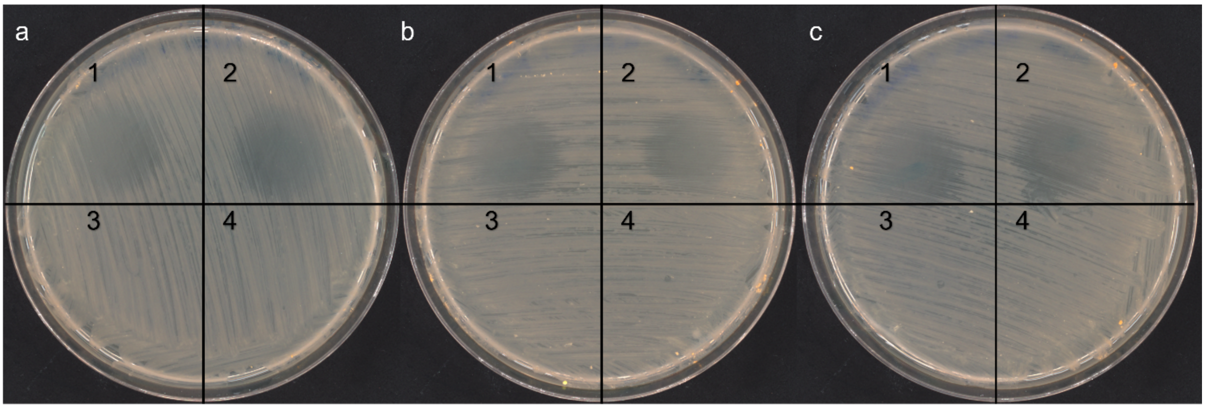

**Figure S5.** Bioactivity of testing of *in vitro* reactions; 1) Gelatinamin. 2) Gelatinamin incubated with AcCoA 3) Gelatinamin incubated with AcCoA and triT 4) triT incubated with AcCoA. All samples are tested in triplicate (a, b and c) against *M. phlei.*

##### 3.2 MS$^2$ of lasso peptides

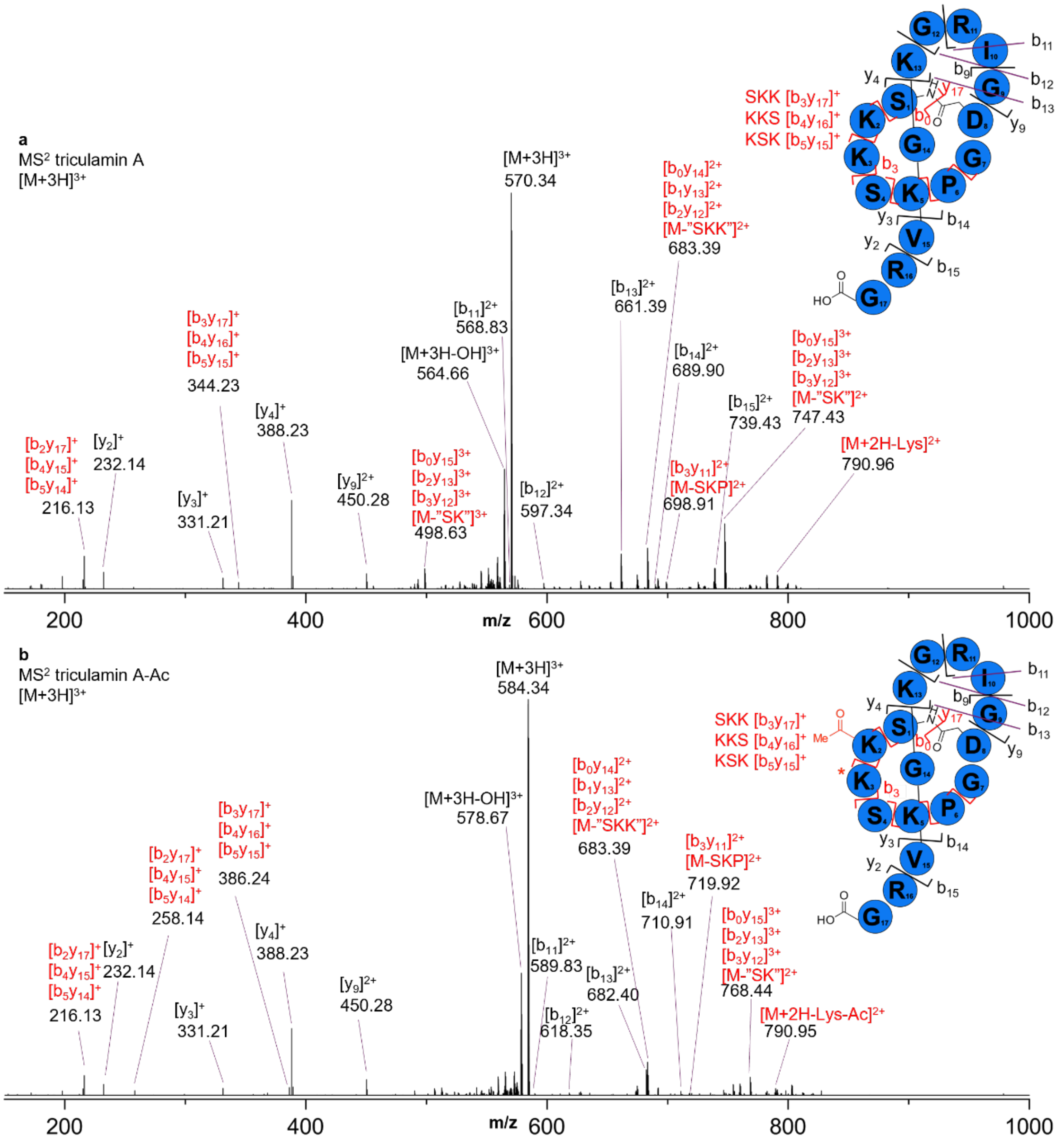

**Figure S6.** MS$^2$ of triculamin A and triculamin A-Ac a) MS$^2$ of the previously NMR-structure resolved triculamin A and the triculamin A-Ac described in this paper. b) MS$^2$ of the $[M+3H]^+$ ion. b and y fragments are in black, whereas double fragments are highlighted in red. Where multiple fragments are possible, they are shown above one another. Fragmentation indicates that the acetylation occurs on position 2-Lys, 3-Lys or 5-Lys.

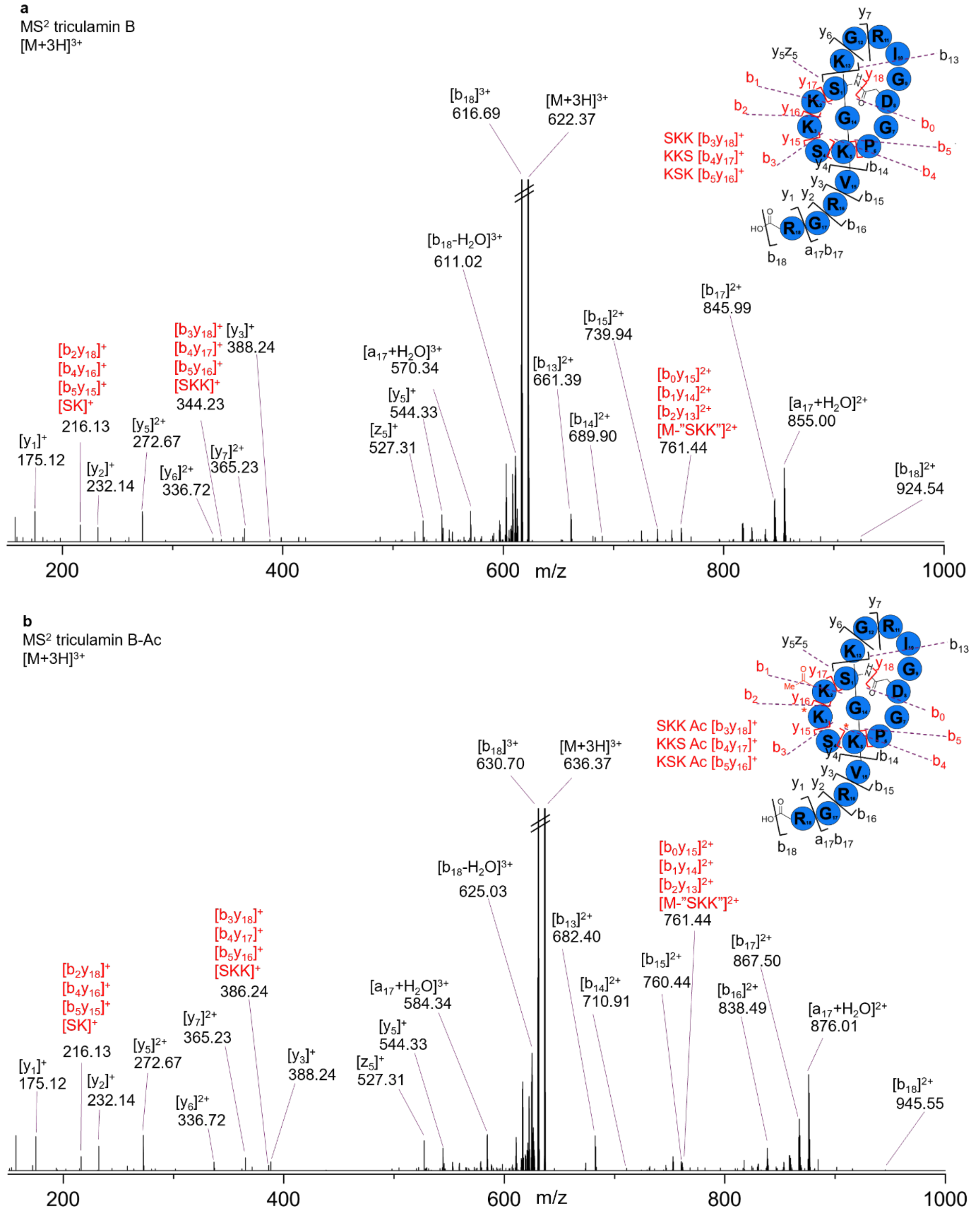

**Figure S7.** MS$^2$ of triculamin B and triculamin B-Ac a) MS$^2$ of the $[M+3H]^+$ ion of triculamin B (a) and triculamin B-Ac (b). b and y fragments are in black, whereas double fragments are highlighted in red. Fragmentation indicates that the acetylation occurs on position K2, K3 or K5.

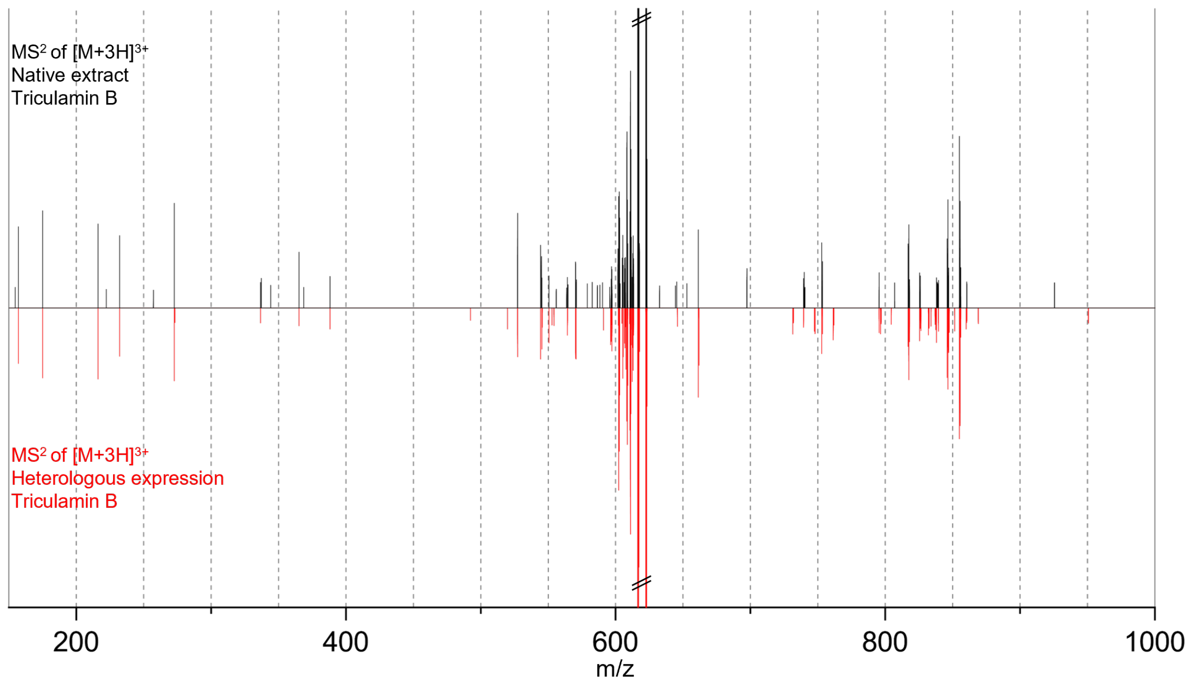

**Figure S8.** The MS$^2$ spectra of triculamin B from either *S. triculaminicus* (black) or *S. albus* heterologous expression (red). The MS$^2$ confirms that the heterologously expressed triculamin B is identical to extracted triculamin B.

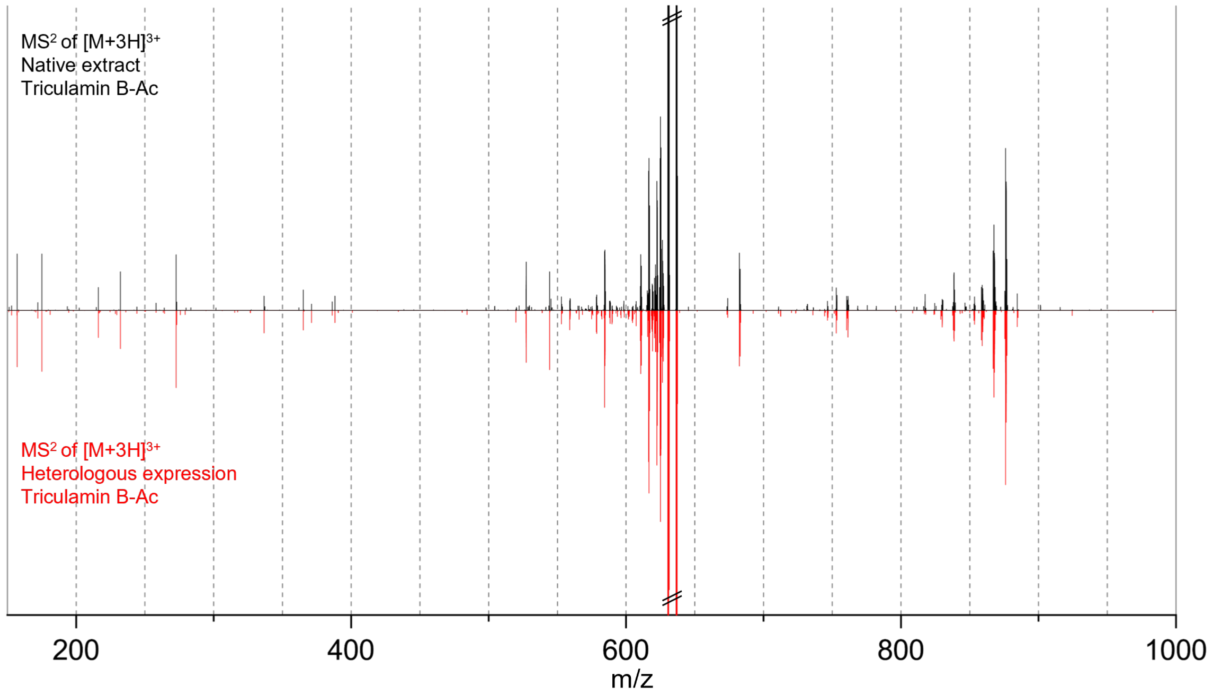

**Figure S9.** The MS$^2$ spectra of triculamin B-Ac from either *S. triculaminicus* (black) or *S. albus* heterologous expression (red). The MS$^2$ confirms that the heterologously expressed triculamin B-Ac is identical to extracted triculamin B-Ac from *S. triculaminicus.*

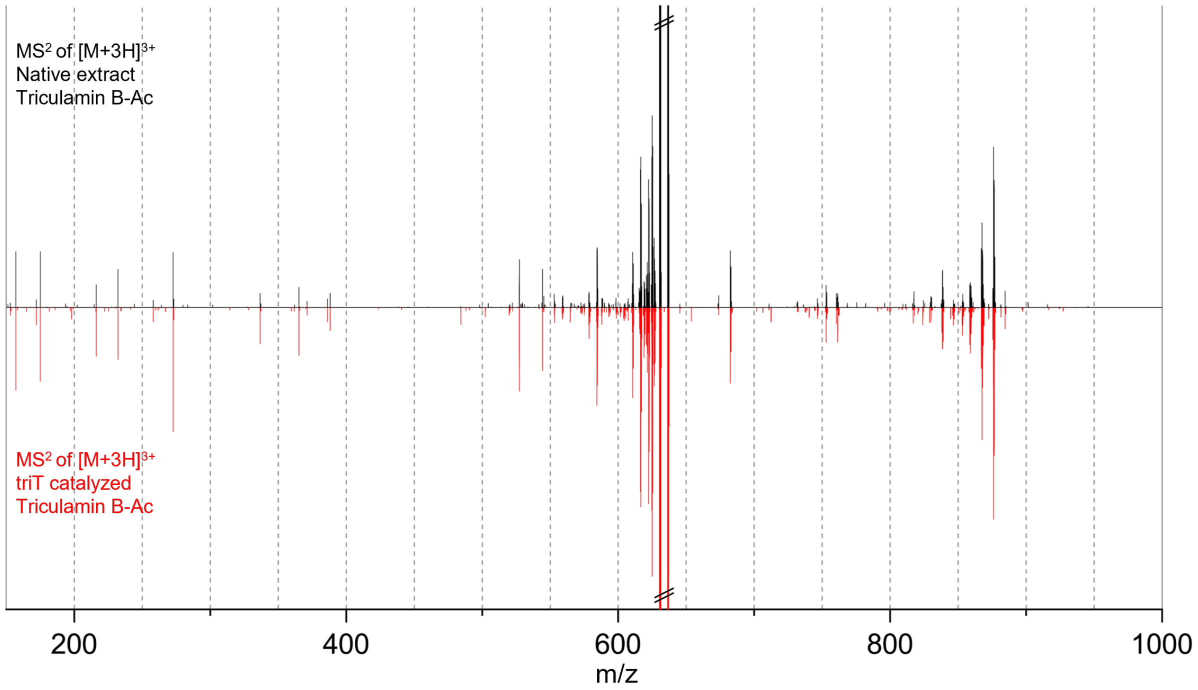

**Figure S10.** $MS^2$ comparison of triculamin B-Ac isolated from *S. triculaminicus* and *in vitro* acetylated triculamin B-Ac. The comparison *in vitro* acetylated triculamin B and triculamin B-Ac extracted from *S. triculaminicus* confirms the same acetylation position.

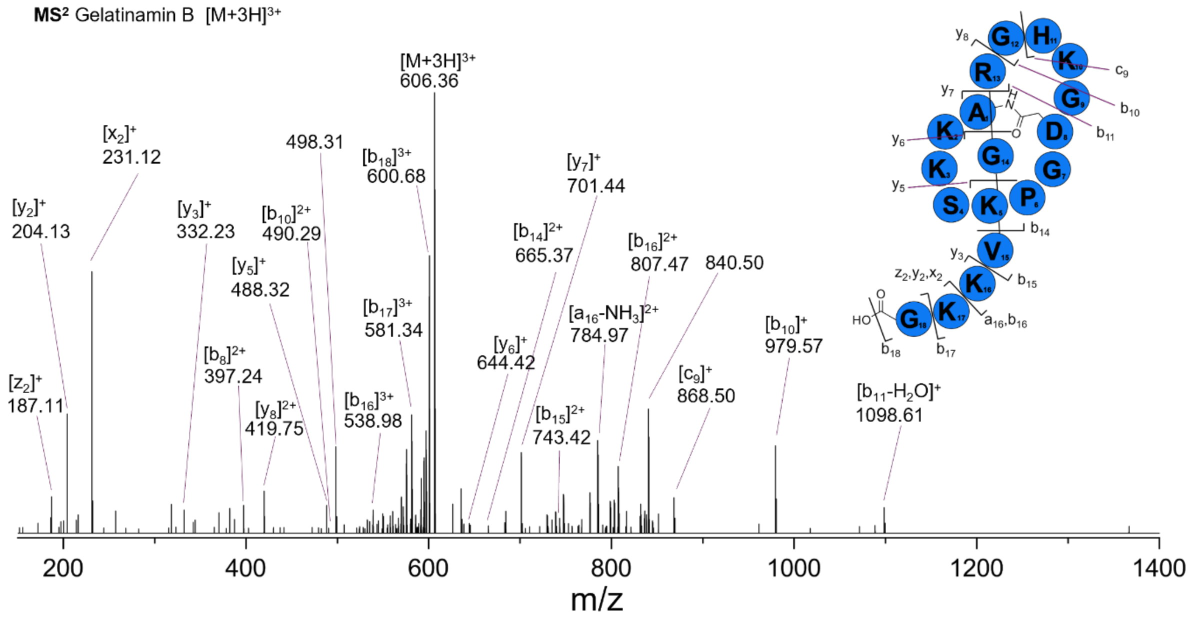

**Figure S11.** $MS^2$ of the $[M+3H]^+$ gelatinamin B. The less abundant gelatinamin B can also be confirmed by the fragmentation pattern.

##### 3.3 Comparison of AlphaFold3 Predicted Lasso Macrocyclases with the Crystal Structure of *E. Coli* Asparagine Synthetase

Lasso macrocyclases all share homology with asparagine synthetases (EC 6.3.5.4), enzymes that catalyze the formation of asparagine from ATP and glutamine.[17] In this reaction, ATP activates the sidechain carboxylic acid of aspartate, while free ammonia, released from glutamine, reacts with the AMP-activated aspartate to yield asparagine.

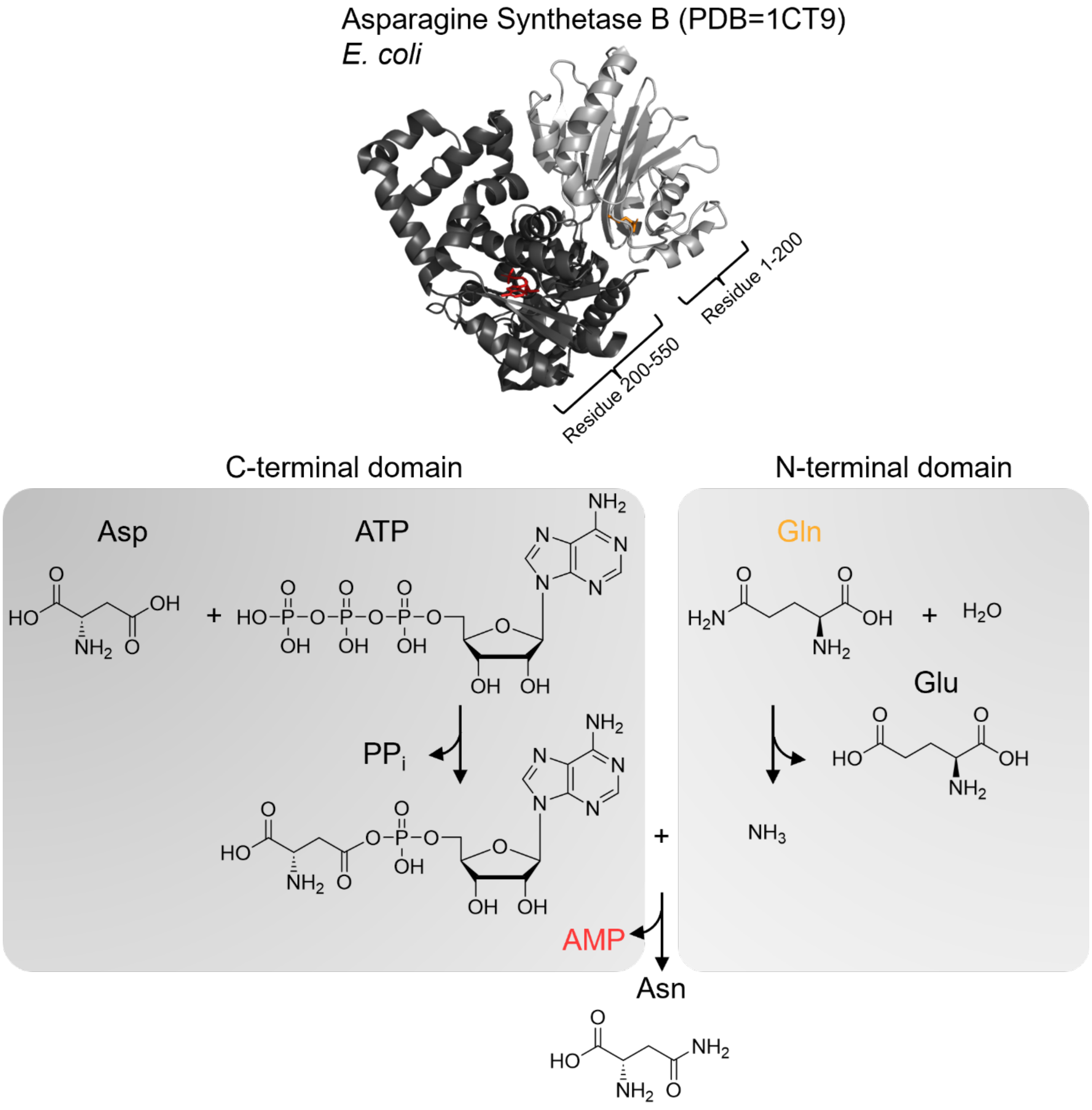

**Figure S12.** Structural representation and catalytic mechanism of the enzyme involved in asparagine biosynthesis. (Top) The crystal structure of the *E. coli* asparagine synthetase, highlighting key regions involved in substrate binding and catalysis. The ATP-binding (AMP) site is shown in red, and the glutamine-binding site is indicated in orange. (Bottom left) The reaction scheme illustrates the initial step, where aspartate (Asp) reacts with ATP, forming an aspartyl-adenylate intermediate and releasing pyrophosphate (PPi). (Bottom right) Glutamine (Gln) is hydrolyzed to glutamate (Glu), generating free ammonia ($NH_3$). The ammonia is then transferred to the aspartyl-adenylate intermediate, producing asparagine (Asn) and releasing AMP.

In contrast, the lasso macrocyclase catalyzes the formation of an isopeptide bond between the N-terminus and an aspartate/glutamate sidechain, removing the need for free ammonia. Few structures of asparagine synthetases have been solved, with only one bacterial asparagine synthetase B structure available (PDB 1CT9) and a related beta-lactam synthetases.[18] Although no structures of lasso macrocyclases have been solved, recent advances with AlphaFold facilitate structural comparisons. All experimentally characterized lasso macrocyclases before the characterization of the triculamin-BGC in the present paper have a predicted N-terminal domain structurally like the glutamine-binding domain, raising intriguing questions about its role. Is this apparently redundant domain essential for structural stability, activity, or does it play a direct role in lasso folding?

Supporting Figure S13 shows the crystal structure of the *E. coli* asparagine synthetase bound to AMP and glutamine. Additional AlphaFold 3 predicted structures include the triculamin macrocyclase (triC), palmamin macrocyclase (palC), and the albusnodin macrocyclase (albC), which is the only other reported lasso peptide with an N-acetyltransferase in its BGC. The structural comparison between Asparagine Synthetase (PDB 1CT9), albC and palC highlight their similar structural features. The C-terminal domain is shared among all four macrocyclases, whereas the N-terminal domain (approx. 200aa) is missing from triC.

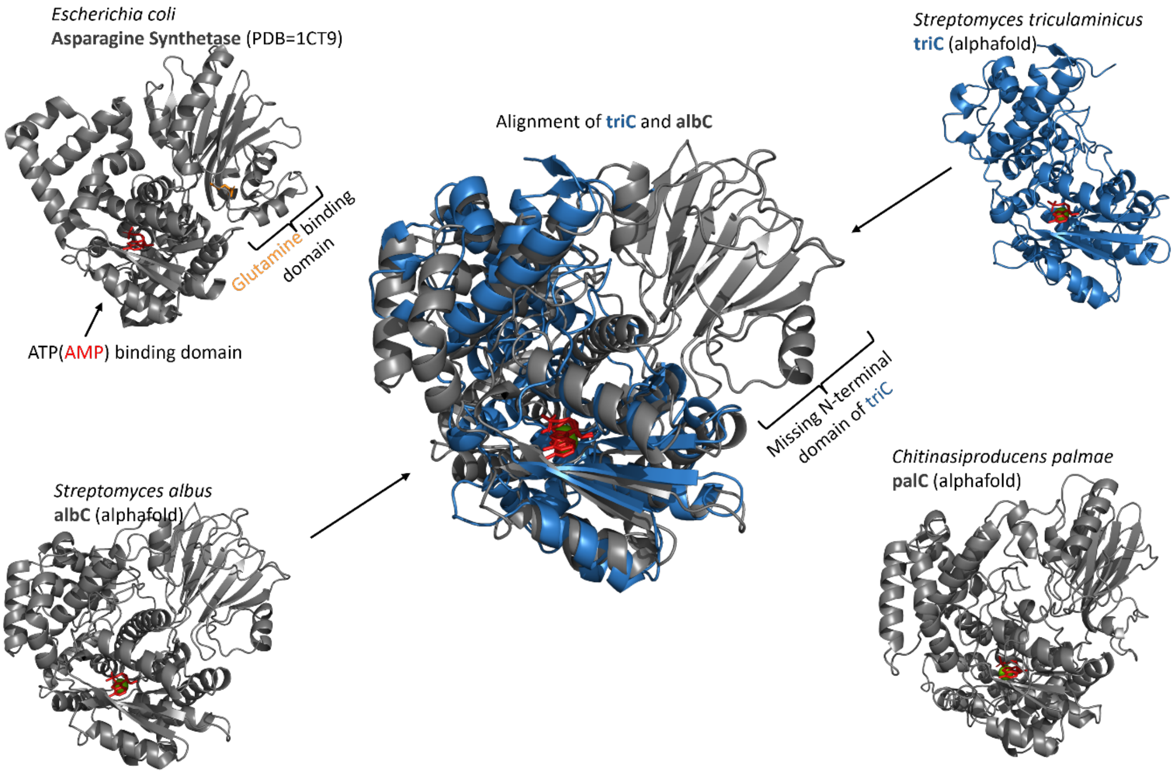

**Figure S13.** Comparison of the crystal structure of asparagine synthetase from *E. coli* with AlphaFold models of macrocyclases from lasso peptide biosynthetic gene clusters (BGCs): albC, triC, and palT. The alignment of triC and albC shows that the two macrocyclases align well, except for an entire N-terminal domain, which is absent in triC.

##### 3.4 Comparison of Acetyltransferase Similarity and Predicted Structures

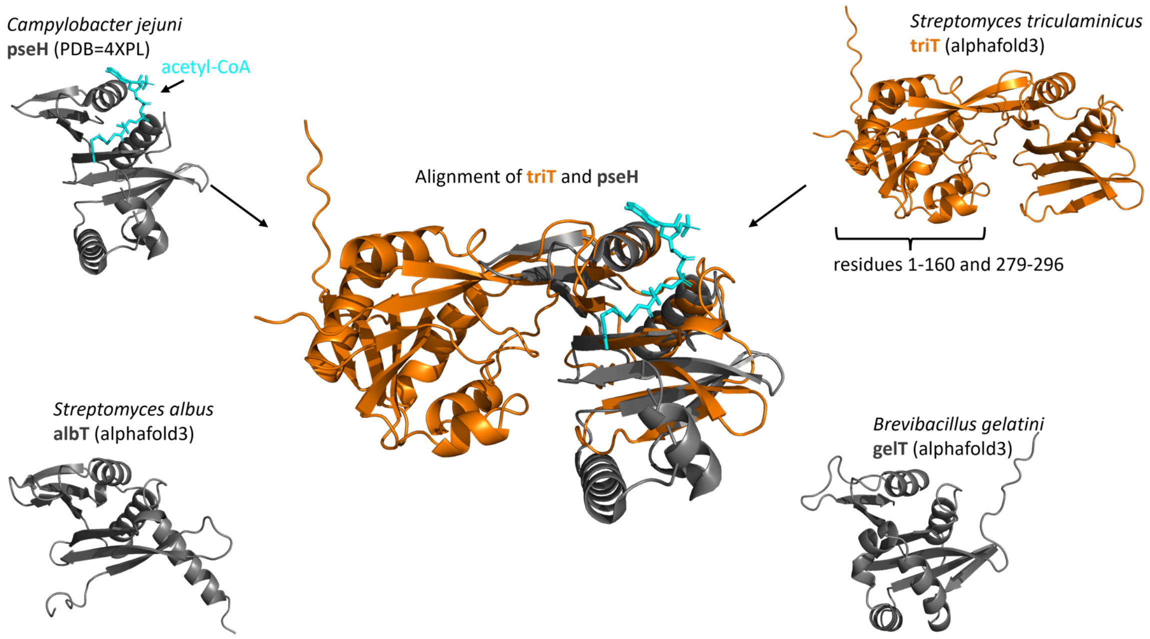

**Figure S14.** Comparison of crystal structure acetyltransferase pseH [PDB:4XPL] with AlphaFold3 structures of acetyltransferases from lasso BGCs (albT, triT and gelT).

##### 3.5 Hypothesized Reaction and Reaction Mechanism of Gelatinamin Clostripain Protease

The *B. gelatini* lasso BGC appeared canonical, containing a precursor (A), RiPP recognition element (B1), leader peptidase (B2), macrocyclase (C), and two ABC transporters (D1 and D2). Based on our findings with triculamin resistance, we were unsurprised to find an N-acetyltransferase within the BGC. However, the presence of gelP, a clostripain (C11-family peptidase), a well-characterized peptidase, was unexpected. Initially, we speculated it might function as a resistance mechanism like the isopeptidases found in some lasso BGCs.[19] Yet, instead of the anticipated gelatinamin, we primarily observed a double-dehydrated form (gelatinamin A). Despite this unanticipated finding, gelatinamin A exhibited extraordinarily similar elution profiles in both cation exchange and reverse-phase chromatography, comparable bioactivity, a similar acetylated and non-acetylated mass pattern, and confirming $MS^2$. Less abundant yet present in sufficient amounts for good $MS^2$ we also characterized the expected lasso peptide (Gelatinamin B) indicating the presence of a tailoring enzyme converting gelatinamin B into gelatinamin A. Reviewing literature about clostripain we became aware of S. Yagisawa *et al* describing the non-native transpeptidase activity of clostripain.[20] This and other reports of other cysteine transpeptidases[21] together with the highly similar transpeptidase activity needed to form gelatinamin A from gelatinamin B, we hypothesized that gelP is responsible for this reaction (Supporting Figure 15a, b). Impressively, the AlphaFold3 predicted structure of gelP in complex with $Ca^{2+}$ and the tail of gelatinamin B (RGVKKG) positions C1 of K17 in direct proximity to the catalytically active Cysteine-193 and Histidine-146 (Supporting Figure 15c). Despite no enzymatic experimental evidence these observations strengthen the hypothesized role of the clostripain family peptidase gelP.

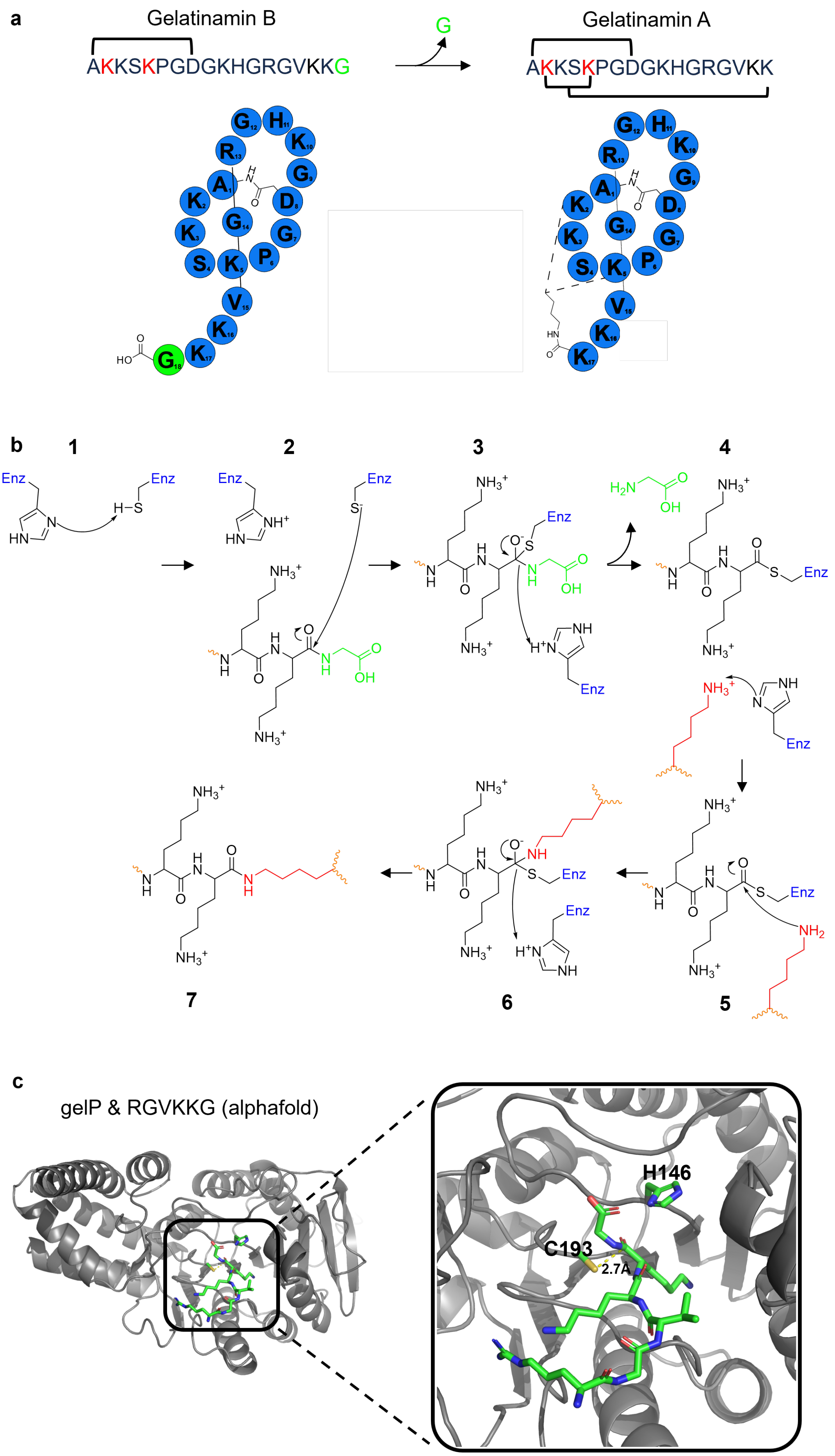
a
Gelatinamin B
AKKSKPGDGKHGRGVKKG
G
Gelatinamin A
AKKSKPGDGKHGRGVKK
b
1
2
3
4
5
6
7
Enz
H2N
OH
c
gelP & RGVKKG (alphafold)
H146
C193
2.7Å

**Figure S15.** Hypothesized function of gelP a) Schematic representation of the double dehydration process of gelatinamin B. b) Proposed reaction mechanism of gelP on gelatinamin B, facilitating the ring closure involving the penultimate N-terminal amino acid and one of the lysines in the ring, resulting in the loss of the N-terminal glycine. c) AlphaFold3 model of gelP in complex with the RGVKKG motif of gelatinamin B, showing the motif folded into the active site.

#### 3.6 Triculamin-like Peptides are Encoded in both Canonical- and Non-canonical BGCs Across Several Bacterial Phyla Supporting

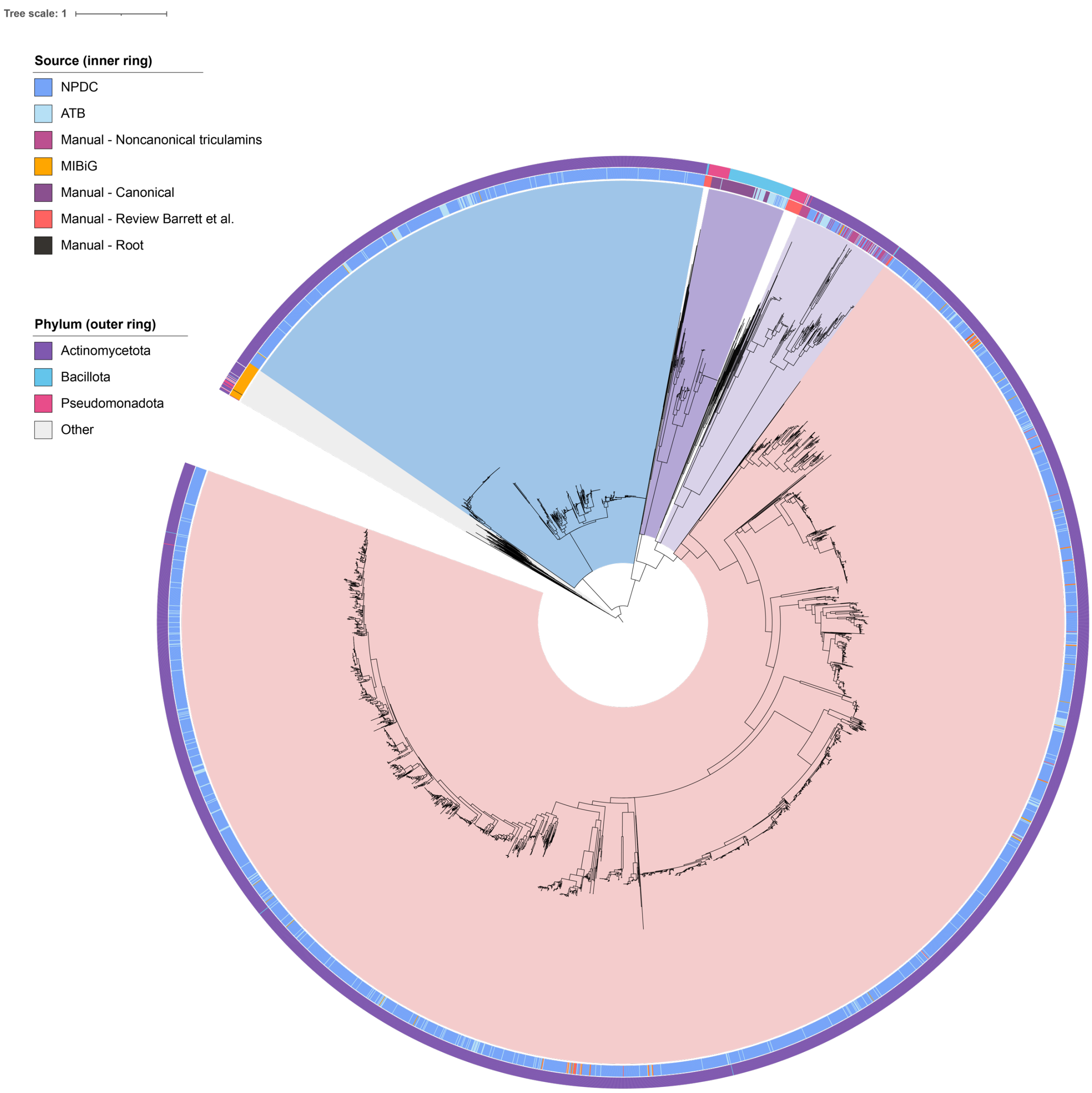

**Figure S16.** Phylogenetic tree of all the macrocyclases used in the study. Clade showing macrocyclase diversity within canonical BGCs encoding triculamin-like lasso peptides (dark purple), clade showing macrocyclase diversity within non-canonical BGCs encoding triculamin-like lasso peptides (light purple), clades showing macrocyclase diversity within previously characterized BGCs from the review by Barrett et al. [1] (white), a clade showing β-lactam synthases (blue), Asn synthases (grey), and a large clade showing “other” macrocyclases (red).

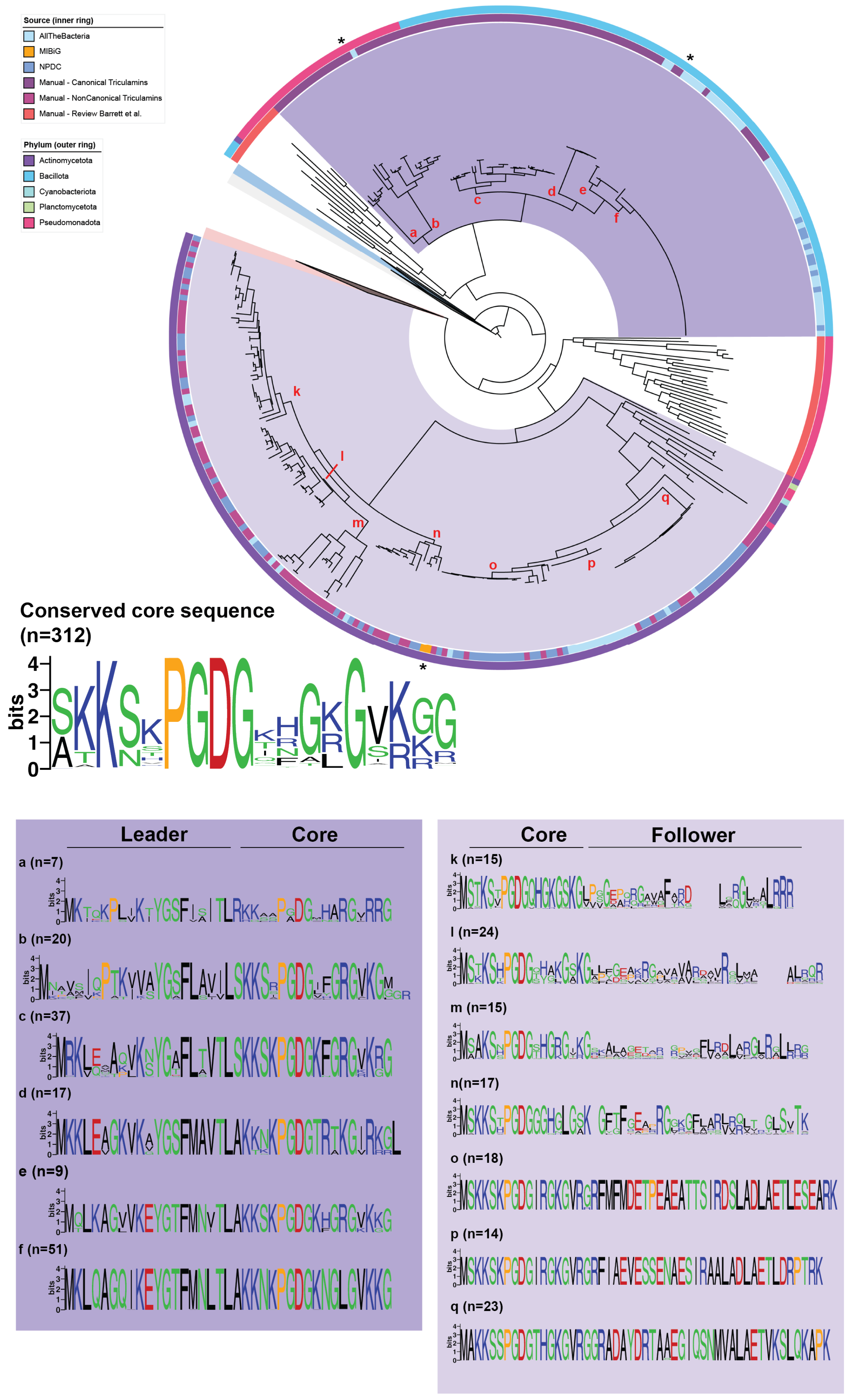

Source (inner ring)
AllTheBacteria
MIBiG
NPDC
Manual - Canonical Triculamins
Manual - NonCanonical Triculamins
Manual - Review Barrett et al.
Phylum (outer ring)
Actinomycetota
Bacillota
Cyanobacteriota
Planctomycetota
Pseudomonadota
Conserved core sequence
(n=312)
bits
Leader
Core
a (n=7)
b (n=20)
c (n=37)
d (n=17)
e (n=9)
f (n=51)
Core
Follower
k (n=15)
l (n=24)
m (n=15)
n(n=17)
o (n=18)
p (n=14)
q (n=23)

**Figure S17.** Phylogenetic tree of the macrocyclases. The tree contains three collapsed notes (grey - Asn synthases, blue - b-lactam synthases, red – other macrocyclase). Clade showing macrocyclase diversity within canonical BGCs encoding triculamin-like lasso peptides (Dark purple), clade showing macrocyclase diversity within non-canonical BGCs encoding triculamin-like lasso peptides (light purple), clades showing macrocyclase diversity within previously characterized BGCs from review by Barrett *et al.*[1] (white). Representations of the core sequence for all triculamin-like peptides as well as the precursors for all subclades has been made using Weblogo[14].

##### 3.7 Observed Variants of Lasso Peptides

Revisiting the *S. triculaminicus* extracts rich in triculamin A, we became aware of multiple triculamin species with varied tail lengths and acetylations. Despite low amounts, these species give insight into the biosynthesis of each lasso peptide described in this paper. Thus, the observed species based on the deconvoluted masses have been compiled in Table S4.

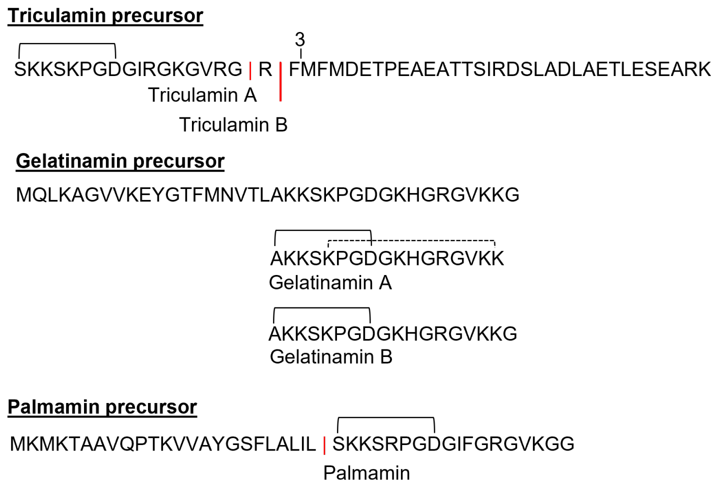

**Figure S18.** Illustration of triculamin, gelatinamin, and palmamin precursors. Lines indicate the ring structures. Triculamin exists in three different variants: ending in Gly (triculamin A), Arg (triculamin B), or Phe. Gelatinamin exists in two variants: with either one ring (gelatinamin B) or two rings (gelatinamin A), the isopeptide bond between the C-terminal and the lysine side chain amine is found on either lys-2 or lys-5. Palmamin was only observed as a single variant.

**Table S4.** The lasso peptides and their variants are named and respective theoretical and observed masses are described. Abundance levels are indicated by ++ (high), +(low) all evaluated by MS traces.

| **Lasso peptide** | **Theoretical mass (Da)** | **Observed mass (Da)** | **Abundance** | ***Source*** |
|---|---|---|---|---|
| Triculamin A | 1707.98563 | 1707.9884 | ++ | *S. triculaminicus* |
| Triculamin A-Ac | 1749.99620 | 1750.0010 | ++ | *S. triculaminicus* |
| Triculamin B | 1864.08675 | 1864.0911 | ++ | *S. triculaminicus* |
| Triculamin B-Ac | 1906.09731 | 1906.1011 | ++ | *S. triculaminicus* |
| Triculamin B-Ac-Ac | 1948.10787 | 1948.0588 | Trace | *S. triculaminicus* |
| Triculamin-3 | 2011.15516 | 2011.1614 | + | *S. triculaminicus* |
| Triculamin-3-Ac | 2053.16572 | 2053.1714 | + | *S. triculaminicus* |
| Triculamin B | 1864.08675 | 1864.0786 | ++ | *S. albus pL99-triACDG* |
| Triculamin B-Ac | 1906.09731 | 1906.0893 | ++ | *S. albus pL99-triACDG* |
| Triculamin B-Ac-Ac | 1948.10787 | 1948.1006 | Trace | *S. albus pL99-triACDG* |
| Gelatinamin A | 1741.02235 | 1741.0124 | ++ | *B. gelatini* |
| Gelatinamin A-Ac | 1783.03292 | 1783.0344 | ++ | *B. gelatini* |
| Gelatinamin A-Ac-Ac | 1825.04348 | 1825.0469 | Trace | *B. gelatini* |
| Gelatinamin-B | 1816.05438 | 1816.0556 | Trace | *B. gelatini* |
| Gelatinamin-B-Ac | 1858.06495 | 1858.0665 | + | *B. gelatini* |
| Unknown | | 1868.9471 | + | *B. gelatini* |
| Unknown | | 1910.9595 | Trace | *B. gelatini* |
| Unknown | | 1797.0163 | Trace | *B. gelatini* |
| Palmamin | 1783.98055 | 1783.9728 | ++ | *B. SP FERM* (pHNF008-palABCD) |

##### 3.8 OSMAC Strategy of Canonical BGC Containing Strains

The OSMAC strategy for the canonical BGCs yielded positive results only for *B. gelatini*, which produced a bioactive compound in both TSA and M9 media (data not shown). An example of this approach is provided in Figure S19 and S20, illustrating a drop assay test where supernatant and pellet samples from *P. tarimensis*, *B. gelatini*, and *C. palmae* were tested. In this instance, the bacteria were cultivated in M9 media, and their bioactivity was evaluated against *M. smegmatis* on MHA.

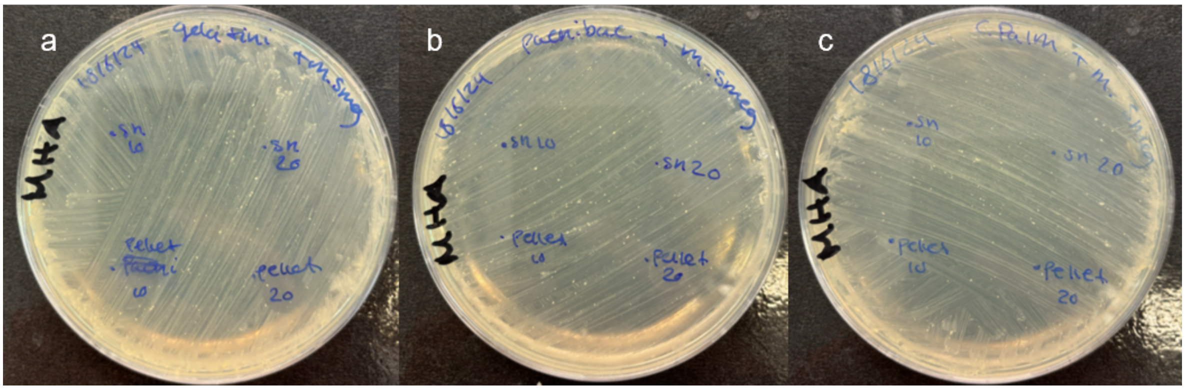

**Figure S19.** Drop assay of supernatant and pellet against *M. smegmatis* from the fermentation of three strains containing canonical triculamin-like BGCs. a) *B. gelatini* drop assay: Left panel shows 10 µL of supernatant (top) and pellet (bottom); right panel shows 20 µL of supernatant (top) and pellet (bottom). b) *P. tarimensis* drop assay: Left panel shows 10 µL of supernatant (top) and pellet (bottom); right panel shows 20 µL of supernatant (top) and pellet (bottom). c) *C. palmae* drop assay: Left panel shows 10 µL of supernatant (top) and pellet (bottom); right panel shows 20 µL of supernatant (top) and pellet (bottom).

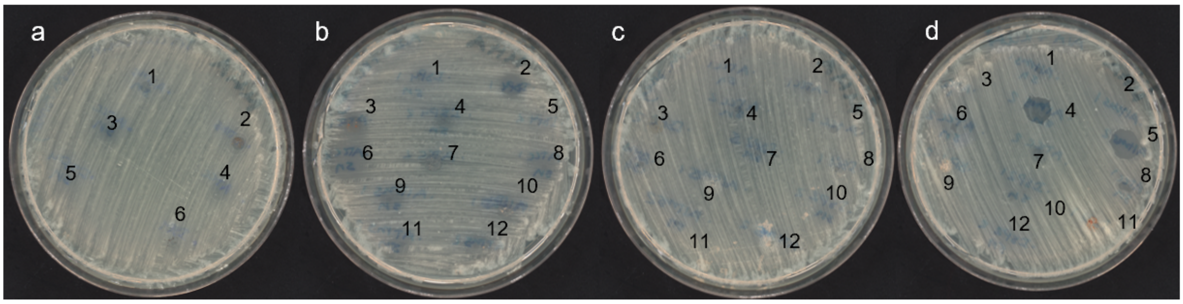

**Figure S20.** Drop assay of supernatant and pellet from the fermentation of four strains containing non-canonical triculamin-like BGCs against *M. smegmatis*. a) *S. triculaminicus* drop assay with single replicates in each condition: (1) supernatant from ISP4, (2) pellet from ISP4, (3) supernatant from ATCC, (4) pellet from ATCC, (5) supernatant from altMS, (6) pellet from altMS. b–d) *S. albus* subsp. *chlorinus*, *S. kurssanovii*, and *S. australiensis* drop assays with duplicate replicates in each condition: (1,2) supernatant and pellet from ISP4, (3,4) supernatant and pellet from ISP4, (5,6) supernatant and pellet from ATCC, (7,8) supernatant and pellet from ATCC, (9,10) supernatant and pellet from altMS, (11,12) supernatant and pellet from altMS.

#### References

[1] S. E. Barrett, D. A. Mitchell, *Trends in Genetics* **2024**, S0168952524001793.

[2] M. R. Green, J. Sambrook, *Cold Spring Harb Protoc* **2020**, *2020*, pdb.prot101196.

[3] H. N. Fernandez, A. M. Kretsch, S. Kunakom, A. E. Kadjo, D. A. Mitchell, A. S. Eustáquio, *ACS Synth. Biol.* **2024**, *13*, 337–350.

[4] F. D. Andersen, K. D. Pedersen, D. Wilkens Juhl, T. Mygind, P. Chopin, E. B. Svenningsen, T. B. Poulsen, M. Braad Lund, A. Schramm, C. H. Gotfredsen, T. Tørring, *J. Nat. Prod.* **2022**, *85*, 1514–1521.

[5] E. W. Sayers, E. E. Bolton, J. R. Brister, K. Canese, J. Chan, D. C. Comeau, R. Connor, K. Funk, C. Kelly, S. Kim, T. Madej, A. Marchler-Bauer, C. Lanczycki, S. Lathrop, Z. Lu, F. Thibaud-Nissen, T. Murphy, L. Phan, Y. Skripchenko, T. Tse, J. Wang, R. Williams, B. W. Trawick, K. D. Pruitt, S. T. Sherry, *Nucleic Acids Research* **2022**, *50*, D20–D26.

[6] M. M. Zdouc, K. Blin, N. L. L. Louwen, J. Navarro, C. Loureiro, C. D. Bader, C. B. Bailey, L. Barra, T. J. Booth, K. A. J. Bozhüyük, J. D. D. Cediel-Becerra, Z. Charlop-Powers, M. G. Chevrette, Y. H. Chooi, P. M. D'Agostino, T. de Rond, E. Del Pup, K. R. Duncan, W. Gu, N. Hanif, E. J. N. Helfrich, M. Jenner, Y. Katsuyama, A. Korenskaia, D. Krug, V. Libis, G. A. Lund, S. Mantri, K. D. Morgan, C. Owen, C.-S. Phan, B. Philmus, Z. L. Reitz, S. L. Robinson, K. S. Singh, R. Teufel, Y. Tong, F. Tugizimana, D. Ulanova, J. M. Winter, C. Aguilar, D. Y. Akiyama, S. A. A. Al-Salihi, M. Alanjary, F. Alberti, G. Aleti, S. A. Alharthi, M. Y. A. Rojo, A. A. Arishi, H. E. Augustijn, N. E. Avalon, J. A. Avelar-Rivas, K. K. Axt, H. B. Barbieri, J. C. J. Barbosa, L. G. Barboza Segato, S. E. Barrett, M. Baunach, C. Beemelmanns, D. Beqaj, T. Berger, J. Bernaldo-Agüero, S. M. Bettenbühl, V. A. Bielinski, F. Biermann, R. M. Borges, R. Borriss, M. Breitenbach, K. M. Bretscher, M. W. Brigham, L. Buedenbender, B. W. Bulcock, C. Cano-Prieto, J. Capela, V. J. Carrion, R. S. Carter, R. Castelo-Branco, G. Castro-Falcón, F. O. Chagas, E. Charria-Girón, A. A. Chaudhri, V. Chaudhry, H. Choi, Y. Choi, R. Choupannejad, J. Chromy, M. S. C. Donahey, J. Collemare, J. A. Connolly, K. E. Creamer, M. Crüsemann, A. A. Cruz, A. Cumsille, J.-F. Dallery, L. C. Damas-Ramos, T. Damiani, M. de Kruijff, B. D. Martín, G. D. Sala, J. Dillen, D. T. Doering, S. R. Dommaraju, S. Durusu, S. Egbert, M. Ellerhorst, B. Faussurier, A. Fetter, M. Feuermann, D. P. Fewer, J. Foldi, A. Frediansyah, E. A. Garza, A. Gavriilidou, A. Gentile, J. Gerke, H. Gerstmans, J. P. Gomez-Escribano, L. A. González-Salazar, N. E. Grayson, C. Greco, J. E. G. Gomez, S. Guerra, S. G. Flores, A. Gurevich, K. Gutiérrez-García, L. Hart, K. Haslinger, B. He, T. Hebra, J. L. Hemmann, H. Hindra, L. Höing, D. C. Holland, J. E. Holme, T. Horch, P. Hrab, J. Hu, T.-H. Huynh, J.-Y. Hwang, R. Iacovelli, D. Iftime, M. Iorio, S. Jayachandran, E. Jeong, J. Jing, J. J. Jung, Y. Kakumu, E. Kalkreuter, K. B. Kang, S. Kang, W. Kim, G. J. Kim, H. Kim, H. U. Kim, M. Klapper, R. A. Koetsier, C. Kollten, Á. T. Kovács, Y. Kriukova, N. Kubach, A. M. Kunjapur, A. K. Kushnareva, A. Kust, J. Lamber, M. Larralde, N. J. Larsen, A. P. Launay, N.-T.-H. Le, S. Lebeer, B. T. Lee, K. Lee, K. L. Lev, S.-M. Li, Y.-X. Li, C. Licona-Cassani, A. Lien, J. Liu, J. A. V. Lopez, N. V. Machushynets, M. I. Macias, T. Mahmud, M. Maleckis, A. M. Martinez-Martinez, Y. Mast, M. F. Maximo, C. M. McBride, R. M. McLellan, K. M. Bhatt, C. Melkonian, A. Merrild, M. Metsä-Ketelä, D. A. Mitchell, A. V. Müller, G.-S. Nguyen, H. T. Nguyen, T. H. J. Niedermeyer, J. H. O'Hare, A. Ossowicki, B. O. Ostash, H. Otani, L. Padva, S. Paliyal, X. Pan, M. Panghal, D. S. Parade, J. Park, J. Parra, M. P. Rubio, H. T. Pham, S. J. Pidot, J. Piel, B. Pourmohsenin, M. Rakhmanov, S. Ramesh, M. H. Rasmussen, A. Rego, R. Reher, A. J. Rice, A. Rigolet, A. Romero-Otero, L. R. Rosas-Becerra, P. Y. Rosiles, A. Rutz, B. Ryu, L.-A. Sahadeo, M. Saldanha, L. Salvi, E. Sánchez-Carvajal, C. Santos-Medellin, N. Sbaraini, S. M. Schoellhorn, C. Schumm, L. Sehnal, N. Selem, A. D. Shah, T. K. Shishido, S. Sieber, V. Silviani, G. Singh, H. Singh, N. Sokolova, E. C. Sonnenschein, M. Sosio, S. T. Sowa, K. Steffen, E. Stegmann, A. B. Streiff, A. Strüder, F. Surup, T. Svenningsen, D. Sweeney, J. Szenei, A. Tagirdzhanov, B. Tan, M. J. Tarnowski, B. R. Terlouw, T. Rey, N. U. Thome, L. R. Torres Ortega, T. Tørring, M. Trindade, A. W. Truman, M. Tvilum, D. W. Udwary, C. Ulbricht, L. Vader, G. P. van Wezel, M. Walmsley, R. Warnasinghe, H. G. Weddeling, A. N. M. Weir, K. Williams, S. E. Williams, T. E. Witte, S. M. W. Rocca, K. Yamada, D. Yang, D. Yang, J. Yu, Z. Zhou, N. Ziemert, L. Zimmer, A. Zimmermann, C. Zimmermann, J. J. J. van der Hooft, R. G. Linington, T. Weber, M. H. Medema, *Nucleic Acids Research* **2024**, gkae1115.

[7] B. Buchfink, K. Reuter, H.-G. Drost, *Nat Methods* **2021**, *18*, 366–368.

[8] M. Hunt, L. Lima, D. Anderson, J. Hawkey, W. Shen, J. Lees, Z. Iqbal, **2024**, DOI 10.1101/2024.03.08.584059.

[9] E. Kalkreuter, S. A. Kautsar, D. Yang, C. D. Bader, C. N. Teijaro, L. L. Fluegel, C. M. Davis, J. R. Simpson, L. Lauterbach, A. D. Steele, C. Gui, S. Meng, G. Li, K. Viehrig, F. Ye, P. Su, A. F. Kiefer, A. Nichols, A. J. Cepeda, W. Yan, B. Fan, Y. Jiang, A. Adhikari, C.-J. Zheng, L. Schuster, T. M. Cowan, M. J. Smanski, M. G. Chevrette, L. P. S. De Carvalho, B. Shen, **2023**, DOI 10.1101/2023.12.14.571759.

[10] K. Katoh, D. M. Standley, *Molecular Biology and Evolution* **2013**, *30*, 772–780.

[11] S. Capella-Gutiérrez, J. M. Silla-Martínez, T. Gabaldón, *Bioinformatics* **2009**, *25*, 1972–1973.

[12] M. N. Price, P. S. Dehal, A. P. Arkin, *PLoS ONE* **2010**, *5*, e9490.

[13] M. A. Larkin, G. Blackshields, N. P. Brown, R. Chenna, P. A. McGettigan, H. McWilliam, F. Valentin, I. M. Wallace, A. Wilm, R. Lopez, J. D. Thompson, T. J. Gibson, D. G. Higgins, *Bioinformatics* **2007**, *23*, 2947–2948.

[14] G. E. Crooks, G. Hon, J.-M. Chandonia, S. E. Brenner, *Genome Res.* **2004**, *14*, 1188–1190.

[15] N. Sun, Z.-B. Wang, H.-P. Wu, X.-M. Mao, Y.-Q. Li, *Ann Microbiol* **2012**, *62*, 1541–1546.

[16] Y. Tong, C. M. Whitford, K. Blin, T. S. Jørgensen, T. Weber, S. Y. Lee, *Nat Protoc* **2020**, *15*, 2470–2502.

[17] T. M. Larsen, S. K. Boehlein, S. M. Schuster, N. G. J. Richards, J. B. Thoden, H. M. Holden, I. Rayment, *Biochemistry* **1999**, *38*, 16146–16157.

[18] M. T. Miller, B. O. Bachmann, C. A. Townsend, A. C. Rosenzweig, *Nat. Struct Biol.* **2001**, *8*, 684–689.

[19] J. R. Chekan, J. D. Koos, C. Zong, M. O. Maksimov, A. J. Link, S. K. Nair, *J. Am. Chem. Soc.* **2016**, *138*, 16452–16458.

[20] S. Yagisawa, S. Watanabe, T. Takaoka, H. Azuma, *Biochemical Journal* **1990**, *266*, 771–775.

[21] Y. Zong, S. K. Mazmanian, O. Schneewind, S. V. L. Narayana, *Structure* **2004**, *12*, 105–112.
